# Resolution of the proteome, transcript and ionome dynamics upon Zn re-supply in Zn-deficient Arabidopsis

**DOI:** 10.1101/600569

**Authors:** Borjana Arsova, Sahand Amini, Maxime Scheepers, Dominique Baiwir, Gabriel Mazzucchelli, Monique Carnol, Bernard Bosman, Patrick Motte, Edwin de Pauw, Michelle Watt, Marc Hanikenne

## Abstract

- Regulation of plant Zn acquisition is poorly understood, while Zn deficiency affects over 2 billion people worldwide. We therefore dissected the dynamic response to changes in Zn supply in Arabidopsis.
- Hydroponically-grown Zn starved plants were re-supplied with Zn. Subsequent time-resolved sampling strategy allowed concomitant quantification of the dynamics of Zn uptake, microsomal and soluble proteins, and specific transcripts, in space (roots and shoots) and time.
- Zn accumulates in roots within 10min, but 8h are needed before shoot Zn increases. By 8h, root Zn concentration was ~60% of non-starved plants. Overexpressed root Zn transporters further peaked in 10-30min post re-supply, before reaching a minimum in 120min and 200 ppm Zn. Zn-responding signaling/regulatory molecules include receptor and MAP kinases, calcium signaling proteins, phosphoinositides, G-proteins, COP9 signalosome members, as well as multiple transcription factors.
- Zn acquisition is a highly controlled dynamic process. Our study identifies novel players in Zn homeostasis and points to cross-talk with other nutrients. It paves the way for directed investigation of so far omitted candidates which dynamically respond to sudden changes in Zn supply but are expressed at similar levels at steady-state Zn deficiency and sufficiency.

## Introduction

Zinc (Zn) is an essential micronutrient for all organisms (Ricachenevsky *et al.*, 2015). In plants, Zn is important for plant growth and development, as well as increased tolerance to abiotic stress and resistance against pathogens (Broadley *et al.*, 2007; Palmgren *et al.*, 2008; Huang *et al.*, 2009; Kodaira *et al.*, 2011). Zn deficiency can cause loss of enzyme activity, photosynthesis inhibition resulting in leaf chlorosis, reduced root growth, poor floral fertility, low biomass, and poor quality in reproductive structures. Zn excess is also detrimental to plants, resulting in reduced growth, chlorosis and altered nutrient homeostasis (Broadley *et al.*, 2007; Nouet *et al.*, 2011; Sinclair & Krämer, 2012). In nature, plants face these opposite challenges. Soils are indeed Zn deficient in large areas worldwide, resulting in about one third of the human population risking Zn deficiency with severe health impact (Alloway, 2009). In contrast, Zn-contaminated soils are the result of a range of anthropogenic activities (Broadley et al. 2007). On individual basis, plant roots may have to face non-homogeneous Zn availability in soils when growing and have to rapidly adjust their response. Thus, plants encounter highly variable conditions in time and space during their lifetime and possess sophisticated molecular mechanisms, referred to as Zn homeostasis, that adjust the amount of Zn in various tissues and during development to a wide range of Zn availability, to ensure optimum nutrition and growth (Palmer & Guerinot, 2009; Briat *et al.*, 2015).

Cellular Zn, as well as copper (Cu), cadmium (Cd) and possibly manganese (Mn), uptake is mediated by Zinc-Regulated Transporter/Iron-Regulated Transporter proteins [ZRT/IRT-like proteins (ZIPs)], which are hypothesized to play an important role in Zn absorption from soil (Krämer *et al.*, 2007; Milner *et al.*, 2013; Ricachenevsky *et al.*, 2015). Arabidopsis possesses 15 ZIP transporter-encoding genes of which *ZIP1* to *ZIP10* are transcriptionally upregulated under Zn deficiency. Some ZIP proteins are localized in the root plasma membrane (e.g. ZIP2); some at the tonoplast (e.g. ZIP1). IRT1 is a plasma membrane transporter responsible for primarily iron (Fe), but also Zn and Cd, uptake from soil (Vert *et al.*, 2002). Its transcript and protein levels are increased under excess Zn (Barberon *et al.*, 2011). *IRT2* has a transcriptional response similar to *IRT1*, while the *IRT3* expression is increased upon Zn deficiency (Talke *et al.*, 2006; Palmer & Guerinot, 2009; Vert *et al.*, 2009). The precise function of most ZIPs remains to be elucidated (Ricachenevsky et al., 2015).

After uptake, the amount of Zn available for translocation to shoots results from the balance between root vacuolar storage and radial transport (Hanikenne & Nouet, 2011; Deinlein *et al.*, 2012; Claus *et al.*, 2013). Storage of excess Zn and Zn translocation to shoots are mediated, among other transporters, by Heavy Metal ATPase (HMA) Zn/Cd pumps (Williams & Mills, 2005; Hanikenne & Baurain, 2014) and by Cation Diffusion Facilitator proteins [CDF, called Metal Transport Proteins (MTPs) in plants] (Sinclair & Krämer, 2012). HMA3 is localized to vacuolar membrane and contributes to Zn, and Cd, storage in vacuoles, accommodating Zn excess (Gravot *et al.*, 2004; Morel *et al.*, 2009). MTP1 and MTP3 also transport Zn into the vacuole, and the two corresponding genes are expressed differentially in distinct tissues. *MTP1* is involved in basal Zn tolerance, whereas the *MTP3* gene is induced upon Zn excess or Fe deficiency in a FIT (FER-Like Iron Deficiency-Induced Transcription Factor)-dependent manner to store excess Zn in roots (Colangelo & Guerinot, 2004; Desbrosses-Fonrouge *et al.*, 2005; Arrivault *et al.*, 2006). HMA4 and HMA2 are plasma membrane pumps and transport Zn from the root pericycle cells to the apoplast in the xylem, playing a key role in Zn translocation to shoots (Hussain *et al.*, 2004; Hanikenne *et al.*, 2008). Unlike *HMA4*, the *HMA2* transcript levels are increased under Zn deficiency and are systemically responding to an unidentified shoot-born signal (Wintz *et al.*, 2003b; Sinclair *et al.*, 2018). The *MTP2* gene, which is strongly responding to Zn deficiency, is contributing together with *HMA2* to Zn translocation to shoots in these conditions (Sinclair *et al.*, 2018).

Zn transport over plasma or vacuolar membranes occurs as a free ion or bound to various ligands/chelators. Intracellular Zn ions are exclusively present as chelates, with for instance histidine, nicotianamine (NA) or organic acids, in the cytosol or are compartmentalized primarily in the vacuole (Sinclair & Krämer, 2012; Sharma *et al.*, 2016).

The mechanisms controlling Zn transporter transcriptional regulation are far less known. In contrast to Fe homeostasis (Thomine & Vert, 2013; Brumbarova *et al.*, 2015; Yan *et al.*, 2016), few transcriptional regulators of Zn homeostasis are described. The ZIP gene regulation is partially explained by the function of bZIP19 and bZIP23, two Zn deficiency-responsive TFs that in addition to most ZIPs also regulate NAS (NA synthase) genes in response to Zn deficiency (Assunção *et al.*, 2010; Inaba *et al.*, 2015). Reversible binding of free Zn to Cys/His-rich motif of these TFs is proposed to regulate their activity (Assunção *et al.*, 2013).

Only very basic knowledge exists about how Zn status is sensed within plant tissues, and the information pathway that results in adjusted Zn transport and overall homeostasis remains unknown. Signaling molecules, e.g. phytohormones (Masood *et al.*, 2012; Fan *et al.*, 2014; Wang *et al.*, 2015) as well as nitric oxide and Reactive Oxygen Species (Chmielowska-Bąk *et al.*, 2014), have been implicated in Cd and Zn excess responses and linked to Mitogen-Activated Protein Kinase (MAPK) cascades and calcium signaling (Luo *et al.*, 2016). However, all of these molecules are biosynthetic products whose pathways need to be initiated by a primary signal.

To identify new players of Zn homeostasis in plants, a number of large-scale studies have focused on transcriptional changes (Wintz *et al.*, 2003a; Becher *et al.*, 2004; Talke *et al.*, 2006; van de Mortel *et al.*, 2006). Only a few proteomic studies of Zn homeostasis exist (Farinati *et al.*, 2009; Fukao *et al.*, 2009; Fukao *et al.*, 2011; Chiapello *et al.*, 2015; Lucini & Bernardo, 2015; Zargar *et al.*, 2015b), analyzing the response to Zn excess/toxicity in Arabidopsis. These studies used plants constitutively grown at high Zn whereas deficiency, as a global problem, was not addressed (Inaba *et al.*, 2015; Zargar *et al.*, 2015a). In all cases, tissues were harvested at a single time point, effectively taking a snapshot of the physiological adjustment to increased Zn levels. Here, we used Zn-starved Arabidopsis plants and monitored their proteome in parallel to ionome and selected transcripts, in short time points post Zn re-supply to reveal early and dynamic responses upon a change in Zn supply in plants.

## Materials and methods

### Plant material cultivation, harvest and phenotyping

*Arabidopsis thaliana* (Col-0) was grown with 8h light per day at 100 µE/m^2^.s, 20°c. Seeds were germinated on plates with half MS medium (Murashige & Skoog, 1962) and 1% sucrose. After two weeks, plants were moved onto hydroponic trays (Araponics, Belgium) in modified Hoagland medium (Nouet et al. 2015) with Zn-sufficient medium (1 µM Zn) for two weeks. Fresh medium was exchanged weekly. Then, medium without Zn was used for three weeks. Before harvest, control medium (1 µM Zn) was re-supplied and roots and shoots were harvested 10, 30, 120 and 480minutes post re-supply (Fig. S1). In addition, roots and shoots from Zn-starved plants and from plants grown under 1 µM Zn during the 3 weeks were harvested as controls; in these cases, the respective medium was last exchanged 8 hours before harvest. Harvest took place in a 2h45 window at day end. Two sets of plants were grown in parallel. Plants for elemental analysis were processed as described (Nouet *et al.*, 2015). Plants for molecular analyses were harvested in liquid nitrogen and stored at −80°C. Three biological replicates (pools of 3-4 plants) were obtained for each time point.

### Quantitative RT-PCR

Total RNA was extracted using the RNeasy Plant Mini kit with on-column DNase treatment (Qiagen). cDNAs were synthesized with 1µg of total RNA by the RevertAid H Minus First Strand cDNA Synthesis kit with Oligo dT (Thermo Scientific). Quantitative PCRs were performed using Mesa Green qPCR MasterMix (Eurogentec) in 384-well plates with a Quantstudio Q5 system (Applied Biosystems) and primers listed in Table S1. Reactions were performed in 3 technical replicates for each biological replicate. Relative gene expression levels were calculated by the 2^-ΔΔCt^ method (Pfaffl *et al.*, 2001) using multiple reference genes (*EF1α, At1g18050*) (Nouet *et al.*, 2015) for normalization with the qBase software (Biogazelle).

### Ionome profiling

Plant material was digested and used to perform elemental analysis using inductively coupled plasma atomic emission spectroscopy (ICP-OES) as described (Nouet *et al.*, 2015).

### Sample preparation for mass-spectrometry

Frozen tissue was re-suspended in cold extraction buffer (100mM Hepes/KOH pH7.5, 250mM Sucrose, 10% w/v Glycerol, 5mM EDTA, 5mM Ascorbic acid, 2% PVPP, 5mM DTT) with protease and phosphatase inhibitors (Sigma Aldrich and Serva). Microsomal fraction was separated as in Kierszniowska et al., (2009).

The microsomal pellet was washed with extraction buffer without PVPP and re-suspended in 100-200µL extraction buffer with SDS. Protein concentration was determined using Bradford (Thermo Fisher Scientific). Where necessary, soluble fraction proteins were pelleted with 3-5 times ice cold acetone. The SDS concentration was adjusted to 1% (w/w). Proteins were reduced with 10mM DTT for 30min, alkylated in 20mM Iodacetamide for 20min in the dark, and reduction was repeated in 21mM final concentration of DTT.

Proteins were precipitated using the 2D Clean-up Kit (GE Healthcare). The pellet was reconstituted in 50mM ammonium bicarbonate and digested with Trypsin (1/50 w/w) overnight at 37°C, followed with a second Trypsin digestion (1/100 w/w) in 80% Acetonitrile at 37°C for 4 hours. The reaction was stopped with 0.5% Trifluoroacetic Acid (w/v). 20µg of digested protein were desalted using 8µg C18 ZipTip (Millipore, Bedford, MA, USA). Approximately 1.5µg protein was injected into Acquity M-Class UPLC (Waters). Peptides were separated using a 180min gradient which increased from 0 to 40% acetonitrile in 150min and then moved up to 80% acetonitrile.

Eluted peptides were analyzed online on a Q Exactive Plus Orbitrap mass spectrometer (Thermo Scientific) operated with nanoESI in positive mode, with TopN-MSMS method, where N = 12. MS spectra were acquired with Mass range - 400 to 1750 *m/z*, Resolution of 70000, AGC target of 1e6 or Maximum injection time of 50 ms. The parameters for MS2 spectra were: Isolation Window of 2.0 *m/z*, Collision energy (NCE) of 25, Resolution of 17500, AGC target of 1e5 or Maximum injection time of 50 ms.

### Proteomics data analysis and candidate selection

The membrane and soluble fractions for both shoot and root tissues for each biological replicate were measured independently. The 72 ‘.raw’ files have been deposited to the ProteomeXchange Consortium via the PRIDE partner repository (Deutsch *et al.*, 2017; Perez-Riverol *et al.*, 2019) with dataset identifiers PXD013049 – root, and PXD013050 - shoot, sample identifiers are in Data S1 (data publicly available after official publication).

Mass spectra were processed with the MaxQuant software v.1.3.0.5 (Cox & Mann, 2008). The following settings were changed from the default: a minimum of 2 peptides for peptide identification out of which one had to be unique, match between runs set to 2min. The proteins were quantified using cRacker (Zauber & Schulze, 2012). After fold change calculations and for MapMan visualization (Usadel *et al.*, 2009), protein intensities were log2 transformed. Biological pathway analysis was performed with MapMan v. 3.6.0RC1 (Usadel *et al.*, 2009). Functional enrichment was performed using the https://usadellab.github.io/MapManJS/test_oo11.html (Schwacke *et al.*, 2019) Data S6, categories were compared to the entire dataset (i.e. all proteins detected in all samples, 6463 proteins).

## Results

### Plant phenotype upon Zn starvation and re-supply

After 3 weeks of Zn starvation, plants had smaller rosettes than control plants, wavy leaf edges and partial chlorosis, which concentrated around the vein regions and the younger leaves in the center of the rosette (Fig. 1a,b). Zn levels in roots and shoots of Zn-starved plants were significantly lower than in Zn-sufficient plants (Fig. 1c).

**Figure 1.**
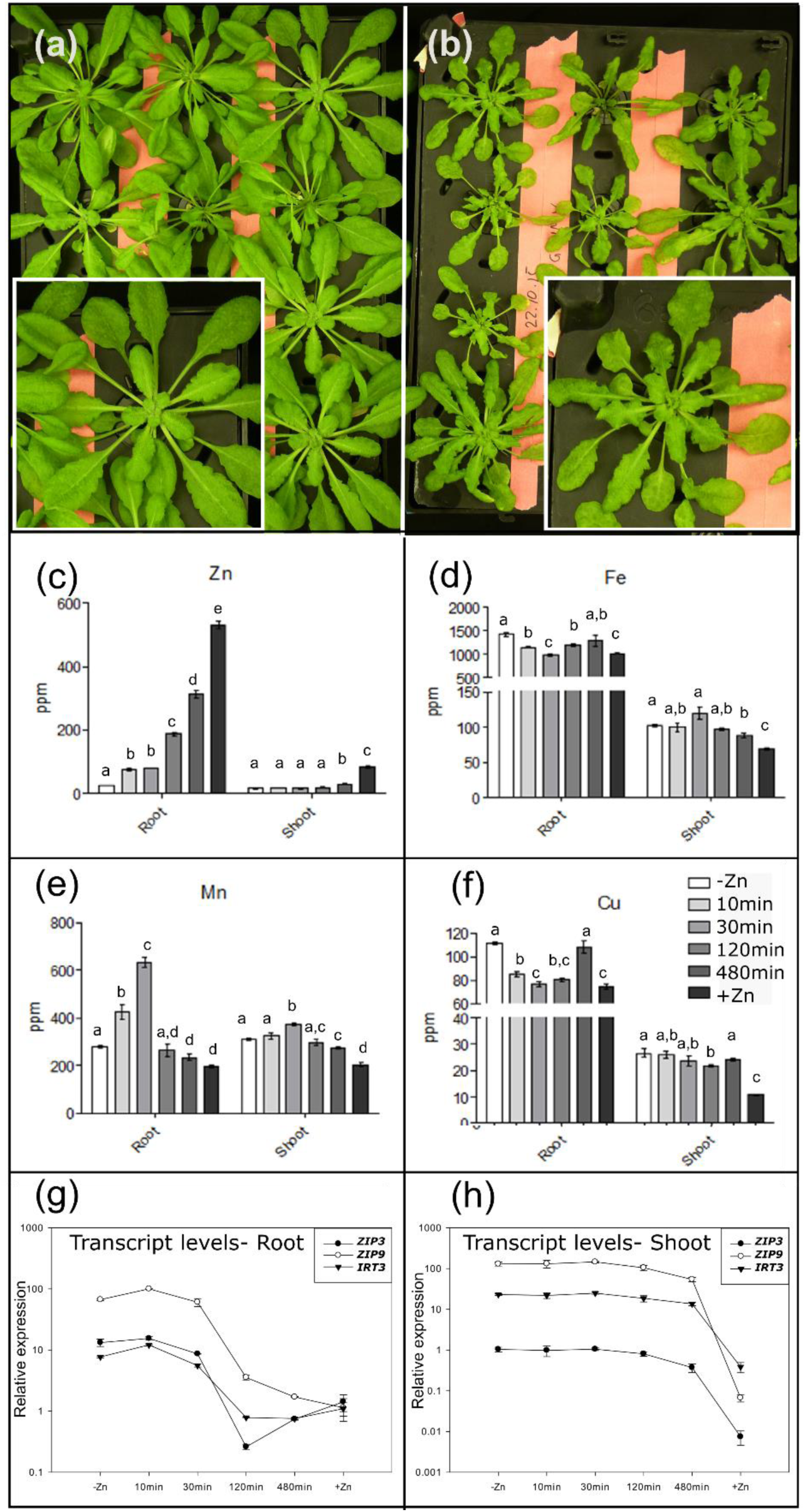
The zinc starvation and re-supply plant material. Plant phenotype of hydroponically grown Arabidopsis plants with sufficient Zn supply (a) and in Zn deficient conditions (b). Elemental analysis of zinc (c), iron (d), manganese (e), copper (f) concentrations for root and shoot tissues throughout the Zn re-supply time series. Gene transcript levels for *ZIP3, ZIP9* and *IRT3* for root (g) and shoot (h) tissues. Legend in (f) applies to bar charts c-f. Error bars show standard deviation (c-f) and standard error (g-h). Each bar/sample point is the average of 3 biological replicates, except root tissues 480 min in plots c-f where n=2. Student’s T-test was performed within each tissue, for data in c-f, where difference with p<0.05 is depicted by the presence of a different letter.

Upon Zn re-supply, the kinetics and magnitude of Zn accumulation in roots was quite striking. Significantly greater Zn was observed in roots of Zn re-supplied plants than of starved plants as early as 10 min upon re-supply (Fig. 1c). This difference remained across most consecutive time points and within 8h Zn concentration in roots increased by 13 fold. Zn accumulation in shoots lagged behind that of roots; a significant increase was not observed until 8h post Zn re-supply. The maximum Zn level reached in shoots was 84 ppm, only 68 ppm higher than the Zn levels in Zn-starved roots, and far from the levels in Zn-sufficient rosettes, indicating that full re-accumulation of Zn would take more than 8h (Fig. 1c).

Fe, Cu and Mn levels were significantly higher in roots and shoots of Zn-starved compared to Zn-sufficient plants (Fig. 1d-f). Fe and Cu levels in roots responded dynamically to Zn re-supply, declining between 10 and 120 min compared to Zn-starved plants followed by re-accumulation. In shoots, only moderate changes in Fe and Cu levels were observed (Fig. 1d,e). In contrast, Mn levels quickly increased in both roots and shoots up to 30min (Fig. 1f) and then decreased to levels observed in Zn-sufficient plants.

### Transcriptional dynamics of known Zn-regulated genes

Transcript levels of known Zn-regulated genes, *ZIP3, ZIP9* and *IRT3* (Talke *et al.*, 2006), were higher in Zn-starved compared to Zn-sufficient plants (Fig. 1g-h), and displayed similar, highly dynamic pattern through time (Fig. 1g-h). In roots, *ZIP9* and *IRT3* transcript levels increased significantly during the first 10min. After 30min, the expression level of the 3 genes decreased compared to Zn-starved conditions to either reach a minimum at 120min, lower than expression in Zn-sufficient plants (*ZIP3* and *IRT3*), or reach the expression level in Zn-sufficient plants (*ZIP9*). *ZIP3* and *IRT3* displayed a sinusoidal-like expression pattern with the maximum and minimum occurring between the starved and sufficient conditions. In shoots, the gene expression levels remained stable until 120min, and only significantly decreased after 480min, remaining much higher than in Zn-sufficient plants (Fig. 1g-h).

In contrast to *ZIP* genes, the *bZIP19* and *bZIP23* transcripts did not show much expression variation, except for *bZIP23* which displayed a slow decrease of expression level through time in shoots, with significantly lower expression in sufficient vs starved conditions (Fig. S2a).

Together with ionome data (Fig. 1c-f), this preliminary expression analysis showed that (i) the plants were indeed Zn-starved at the beginning of the time series, and (ii) the transcript response differed between roots and shoots with, among other, a very rapid response in roots and a delayed response in shoots (Fig. 1g-h).

### The Zn starvation re-supply proteomics dataset

A total of 5249 and 4698 proteins were quantified in roots and shoots, respectively (Fig. 2a), submitted to functional annotation (Fig. 2b) and then examined to detect over- and under-representation of functional categories (Fig. 3, Fig. S3, Data S3, Data S4).

**Figure 2.**
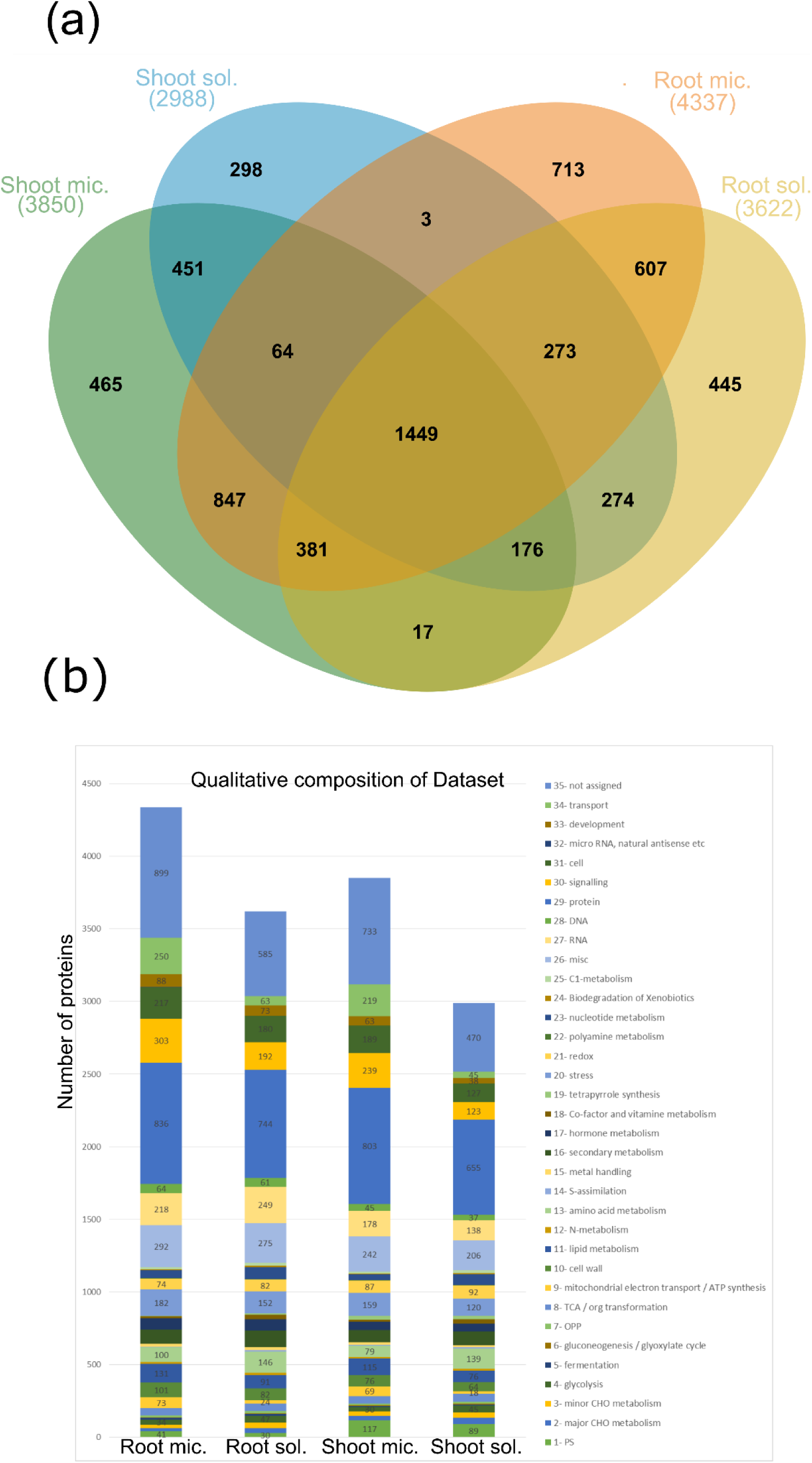
Quantified proteins in root and shoot microsomal and soluble fractions encompassing all six time points of the Zn deficiency and re-supply time series. Overlap of the quantified proteins in each tissue and fraction (a). Qualitative composition of the quantified proteins in each fraction, according to MapMan functional categories (b). The Venn diagram was created using jvenn (Bardou et al, 2014). Microsomal fraction (mic.), soluble fraction (sol.). Total number of proteins for each tissue and fraction listed are in brackets.

**Figure 3.**
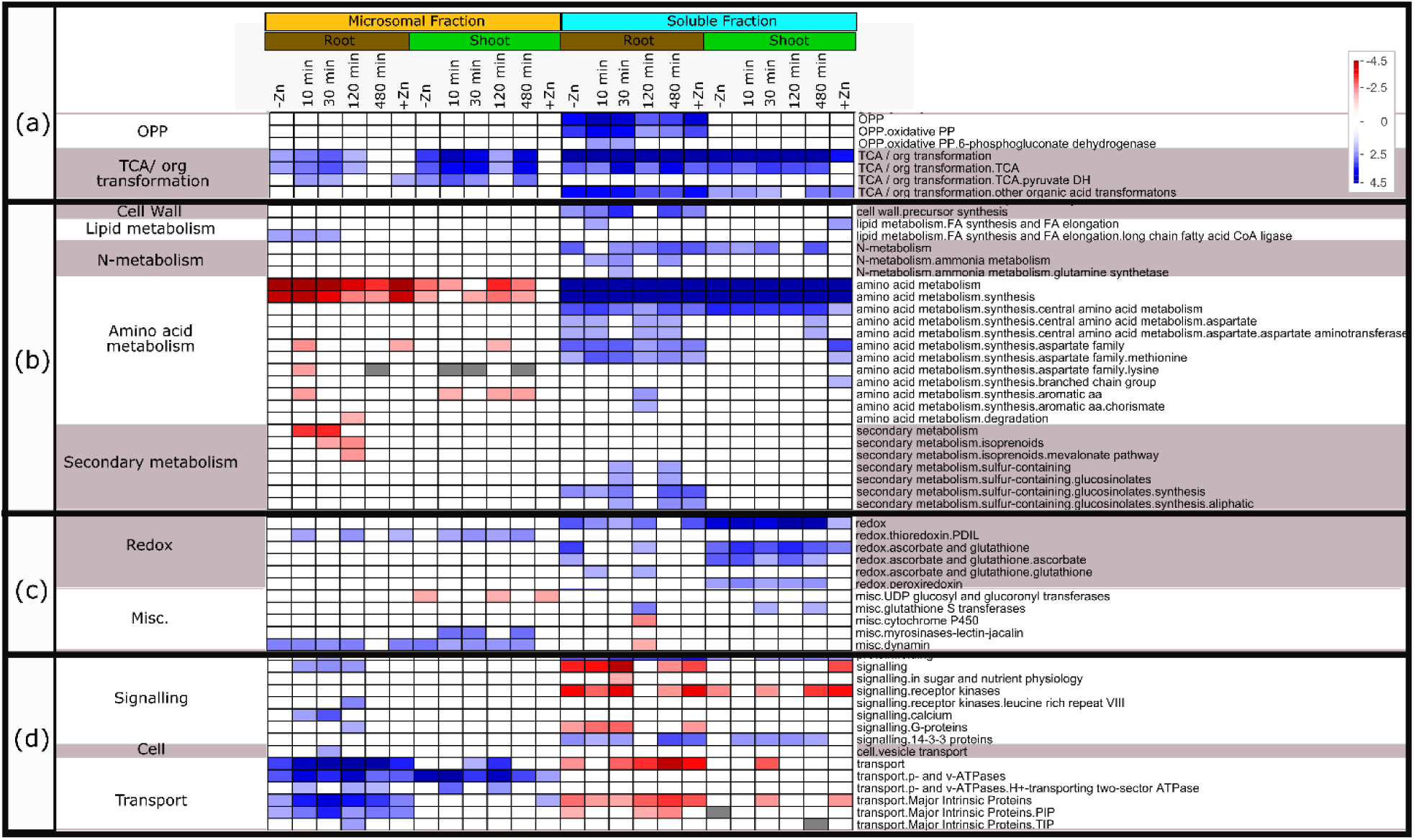
Selected MapMan overview of the root and shoot proteome dataset. A full representation is available as Fig. S3. The protein levels were log_2_-transformed and sorted into MapMan functional categories (Usadel et al., 2009). The protein categories were subjected to a bin-wise Wilcoxon test (ora cutoff = 1.0) and Benjamini-Hochberg multiple testing correction. Data is presented using PageMan (Usadel et al., 2006), with over-represented (blue) and under-represented (red) functional categories, with color scale in the top right corner. OPP: oxidative phosphorylation; TCA: tricarboxylic acid; Misc: Miscellaneous.

#### Soluble fraction

In the soluble fraction, overrepresented functions included photosynthesis, primary energy metabolism (carbohydrate, TCA cycle, glycolysis), amino acid metabolism and proteins involved in redox regulation and protein degradation (Fig. S3). Root and shoot tissues were distinguished based on the representation of the photosynthesis and related functions (Fig. 2b, S3). In roots, proteins involved in cell-wall precursor synthesis were also overrepresented showing a rapid dynamics throughout the time series, whereas nitrogen (N) metabolism responded in both tissues (Fig. 3b). Protein folding and 14-3-3 signaling proteins were overrepresented in both root and shoot tissues, whereas a dynamic response of proteins involved in ubiquitination was observed in roots. Underrepresented functions included RNA processing, protein synthesis, signaling and transport (Fig. S3).

#### Microsomal Fraction

Generally, the membrane enrichment procedure facilitated the identification of a number of transporters and signaling proteins in both roots and shoots that may otherwise have gone undetected due to their low abundance (Fig. 2b, 3d, S3). The shoot microsomal fractions displayed typical over-representation of prokaryotic ribosomal subunits (chloroplast) and proteins involved in photosystem II compared to the root fractions (Fig. S3).

In roots, the transport functional category, including a number of transporter families (Major Intrinsic Proteins, p- and v- ATPases), was enriched and displayed a dynamic response through time (Fig. 3d). Similarly, the signaling category was also highly enriched at 10 to 120min, including calcium signaling molecules and receptor kinases. Enrichment in lipid metabolism, dynamin and vesicle transport were also observed at early time points (Fig. S3). Altogether, this suggested very rapid signaling response to Zn re-supply, combined with dynamic changes in plasma membrane, protein movement to/from the membranes and transport. In the shoot microsomal fraction, further evidence of regulation of protein association to membranes (dynamin, myristoylation) was observed (Fig. 3c, ‘misc’), as well as an increase in p- and v-type ATPase transport proteins (Fig. 3d).

Altogether, this analysis provided an overview of the systemic response to the treatment and revealed that a number of processes responded dynamically to Zn re-supply.

#### Elucidation of time-related dynamic responses

To identify novel players responsible for Zn signaling and regulation of Zn homeostasis, a comparison of each time point post re-supply (i) to Zn deficiency, and (ii) to the previous time point in the time series was conducted, focusing on proteins showing at least a 4-fold change and an adjusted p<0.05 (Fig. S4, Data S5). The rationale for this double comparison was: it identified (i) proteins with a rapid response between two consecutive time points, as well as (ii) proteins that take longer to respond (when a time-point is compared to Zn deficiency). Due to protein extraction and sample preparation procedures, comparisons were always conducted within one tissue and fraction (Data S5) but summarized together for clarity (Data S6).

The dynamic Zn response mobilized the regulation of 1877 proteins from a large set of functional categories (Fig. 4). The number of regulated proteins, and accordingly represented functional categories, was higher in roots than in shoots, with a delayed response in shoots.

**Figure 4.**
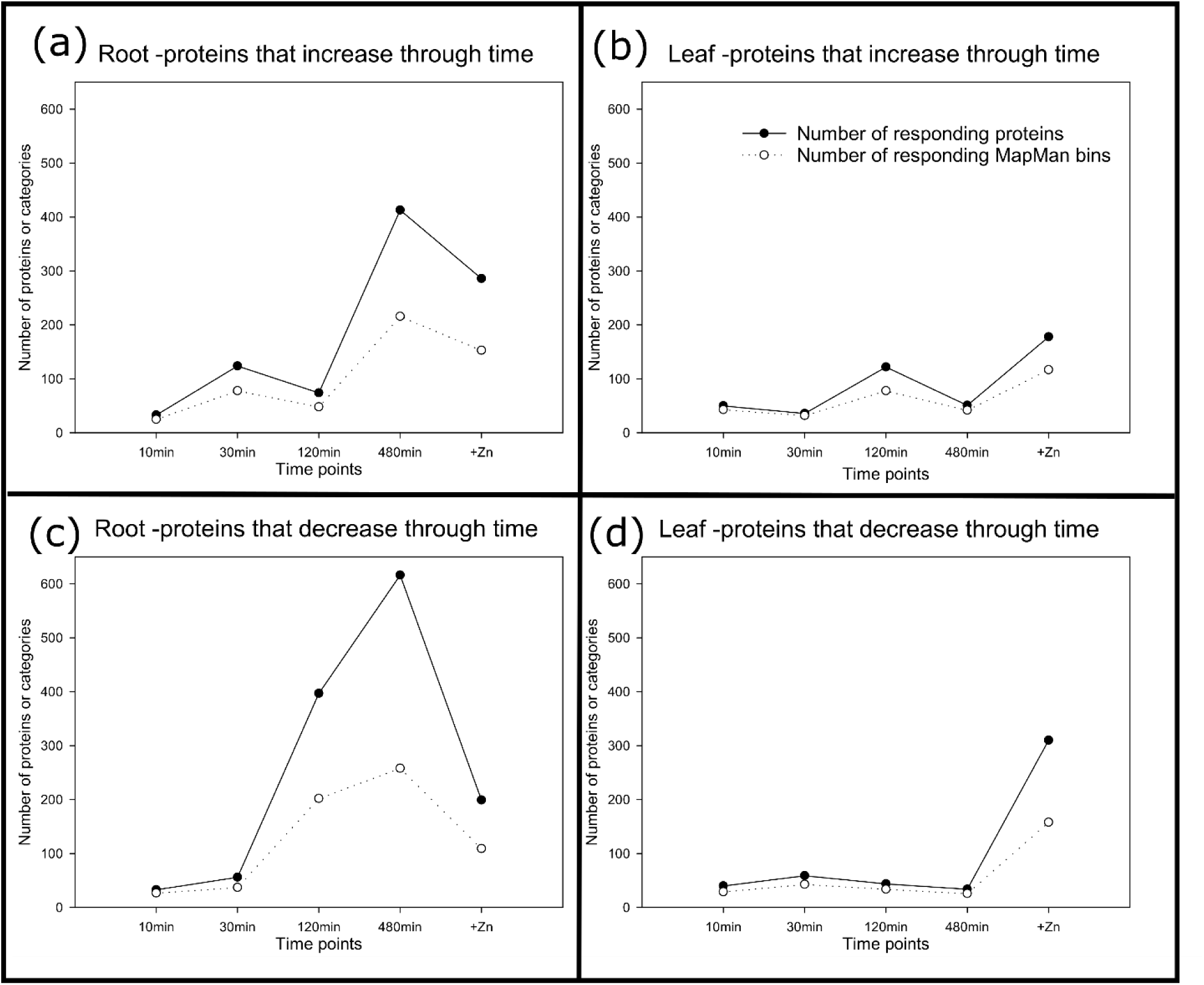
Overview of the up- and down-regulated proteins for the root and shoot datasets during the Zn re-supply time series and the number of unique MapMan (sub)-functional categories represented at each time point. The criteria for protein selection are 4 fold change and a p<0.05 in multiple testing correction as represented in Data S6. Each time point includes the proteins that pass the set thresholds in comparison to the Zn deficiency condition (-Zn) and the previous point in the time-series.

In roots, the number of proteins increasing or decreasing (Fig. 4a, c) through time changed from a few dozens to hundreds, to reach a maximum at 480min indicating that the response was still progressing at this point. At all time-points except 30min, the number of proteins with decreased expression was higher than those with increased expression (e.g. 400 and 72 down- and up-regulated proteins at 120min, respectively), suggesting that rapid dynamic changes are mostly enabled by repression rather than synthesis of new proteins (Fig. 4a, c). In early time points, a limited number of functional categories were involved and included RNA regulation, metal handling, transport, signaling and biotic responses (PR proteins/defensins), highlighting important contribution of those processes to the early response to Zn supply (Fig. 5a,c, Table S2). After 480min, the response mobilized 16 Mapman main functional categories (and more than 200 sub-categories), e.g. transport, signaling, TCA/Organic transformation, cell wall precursor synthesis, peroxidases, ribosomal protein synthesis, central amino acid metabolism, N-metabolism, glycolysis and RNA, thus interlinking Zn acquisition with a large part of the metabolic network (Table S2). It appears that proteins important for plant maintenance under Zn deficient conditions (e.g. stress response, Zn scavenging) remain highly expressed for at least 120min, when Zn levels in roots reached about a third of the Zn found under Zn sufficient conditions (Fig. 1c). Additionally, every functional category that shows proteins decreasing in expression at 120min in roots sees an equal or larger number of proteins decreasing in expression at 480min (Fig. 5c). This indicated a coordinated decrease of proteins belonging to several functional groups, e.g. transport, RNA, signaling, which correlated with Zn influx in the plant (Fig. 1c, 5c).

**Figure 5.**
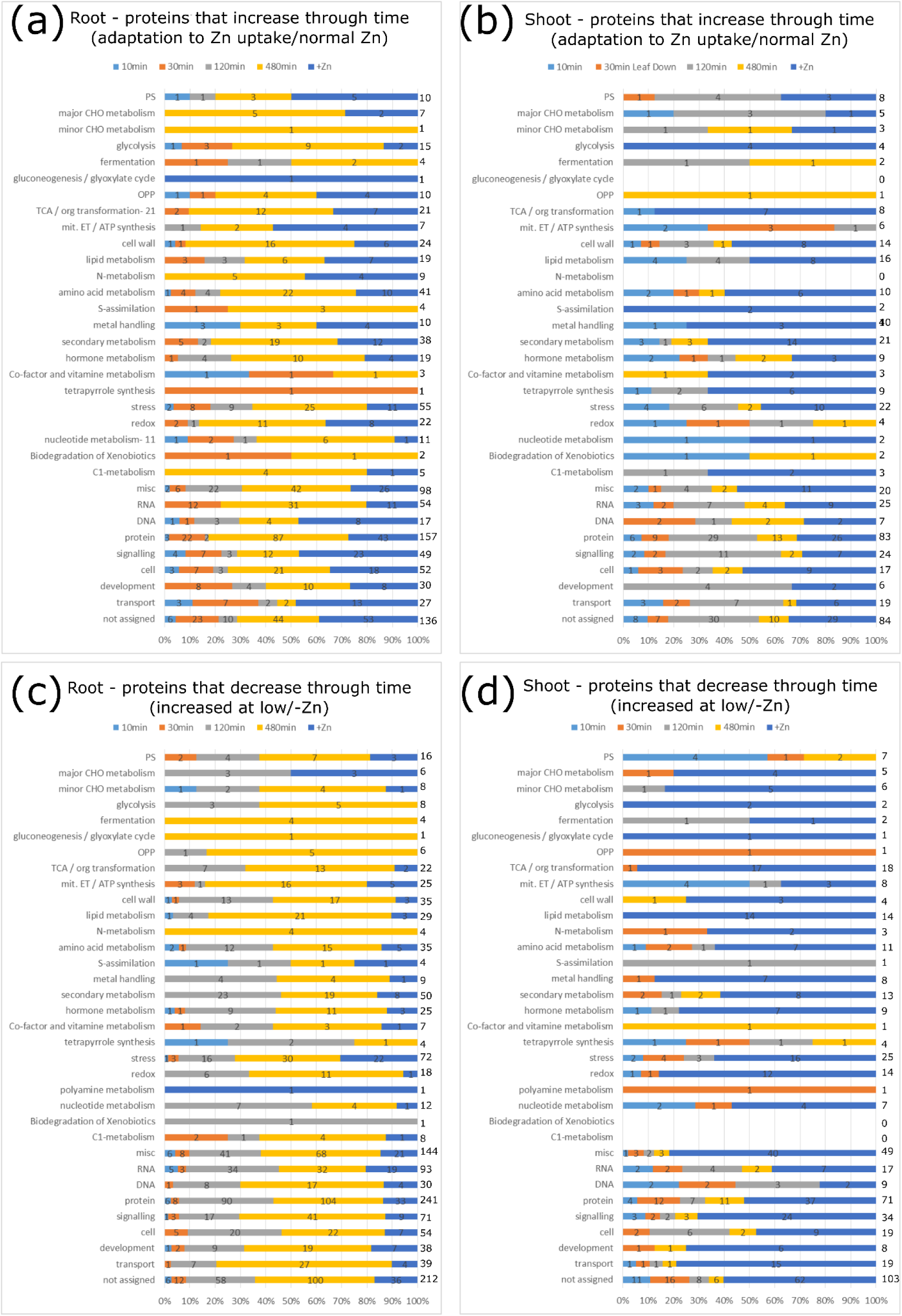
Qualitative composition of the dynamic response after Zn re-supply presented using the MapMan functional categories. Proteins that increase through time in roots (a); Proteins that increase through time in shoots (b); Proteins that decrease through time in roots (c); Proteins that decrease through time in shoots (d). The response for each time point is outlined in colored bars: 10min- light blue, 30min- orange, 120min- grey, 480min- yellow, +Zn- dark blue. Individual AGI can be found in Data S6, information about functional enrichment can be found in Data S6 and precise fold changes can be found in Data S5.

Interestingly, the pattern of up-regulated proteins in shoots displayed a similar dynamics as in roots, but with a delay of one time point (Fig. 4a,b). The first peak of up-regulated proteins (Fig. 4b) appeared after 120min, which precedes the first time point where a significant Zn increase was measured in shoots (Fig. 1c) and included enrichment of signaling proteins, protein synthesis, stress and Calcium transport (Fig. 5b, Table S3). Compared to roots, it also appears that a number of metabolic functions only became activated at 120 and 480min in shoots, whereas these proteins responded at earlier time points in roots (e.g. fermentation, C1 metabolism or oxidative phosphorylation, Fig. 5a,b).

In agreement with the delayed accumulation of Zn in shoots (Fig. 1c), only a small number of proteins were downregulated in response to Zn re-supply, with major difference only observed between Zn deficiency and Zn sufficiency (Fig. 4d), including TCA/org transformation, protein, metal handling, and signaling functions (Fig. 5d, Table S3). Changes of metal handling proteins (ferritins and copper binding proteins) can be linked to lower Fe and Cu concentrations in Zn-sufficient conditions (Fig. 1d,f). It is likely that a longer time series would be required to discover more Zn-responding proteins in shoots, as the second peak observed in roots (480min) is missing in the shoot response curve (Fig. 4a,b).

### Temporal regulation of Zn-related proteins and Zn amount required for recovery from Zn starvation

A number of metal transporters and proteins involved in metal homeostasis were present in the dataset. In agreement with transcriptional regulation (Fig. 1g,h), protein levels of ZIP3, ZIP9 and IRT3 were dramatically increased in both roots and shoots upon Zn deficiency. A similar pattern was observed for putative Zn-transporting proteins or Zn chelator synthesis proteins, including ZIP4, ZIP5, MTP2, HMA2, NAS4 (Fig. 6, Data S2). Most of these proteins were not detected in Zn sufficient conditions, consistent with very low transcript levels. In roots, the protein levels of the 5 ZIP proteins, as well as HMA2 and NAS4, increased moderately in the early time points, with varying amplitude and time dependency, before decreasing at later time. The observed increase in protein levels was slightly delayed compared to transcript regulation, suggesting rapid *de novo* protein synthesis from newly synthesized transcripts. In contrast, the MTP2 protein levels decreased very rapidly in roots upon Zn re-supply. In shoots, most protein levels showed little variation and remained high until 480min, similar to transcript levels (Figure 6). An exception was IRT3 (Fig. 6e) which increased and displayed a maximum at 30min.

**Figure 6.**
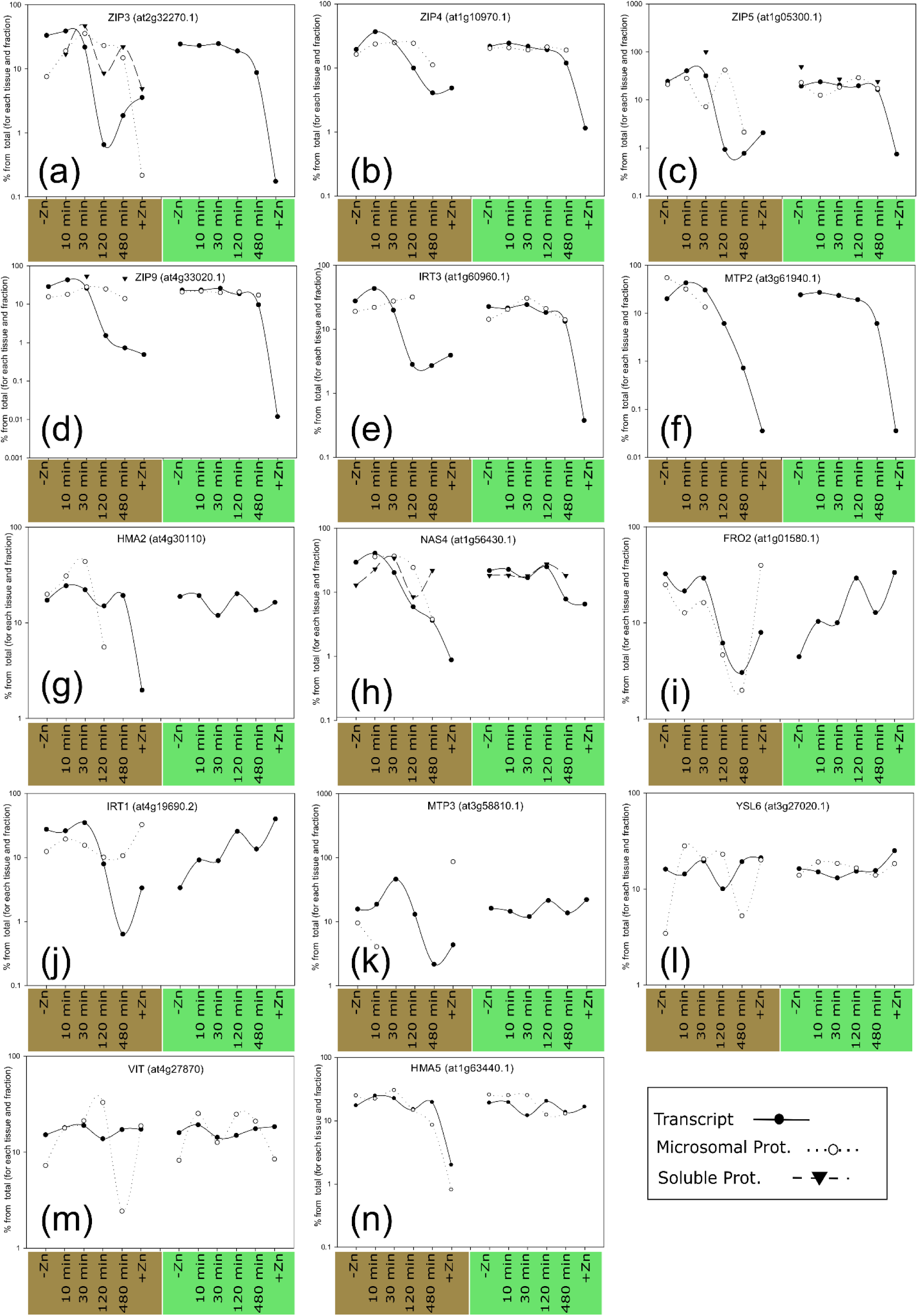
Comparison of transcript and protein regulation profiles for a selection of genes during the Zn starvation and re-supply time series. To present both transcript and protein levels for a gene on the same scale, the transcript expression/protein intensity is presented as the percent for the respective time point from the total of all time points in that tissue and fraction. The reader is reminded that these are relative values to start with and that transcript and protein intensities from each protein fraction are obtained in separate extraction steps. Relative protein intensity was obtained from cRacker (Zauber and Schulze, 2012), transcript levels are relative to *EF1α* and *At1g58050*, the value on the y-axis is common log, brown x-axis labels: root data, green x-axis labels: shoot data.

Next, the relation between Zn and Mn concentrations in tissues was examined in correlation to metal-transporter transcript expression. In both root and shoot comparisons (Fig. 7, 8), the time series was preserved along an axis defined by Zn concentration. In roots, Zn levels negatively correlated with transcript levels, with exception of *HMA2*, and this correlation was not linear but rather followed a quadratic equation with a parabola shape (Fig. 7). Note that the 10 and/or 30min points often appeared as outliers indicating that, at these times, increased transcript levels were associated with increased Zn accumulation. In roots, transcript levels decreased to Zn sufficient levels when Zn amount approximated 200 ppm, which was achieved in 120min. In shoots, and despite a delayed Zn accumulation (Fig. 1c), a similar correlation, mainly influenced by the 480min and Zn-sufficient conditions, was observed indicating that even minute amounts of Zn were sufficient to decrease Zn transporter transcripts (Fig. 8).

**Figure 7.**
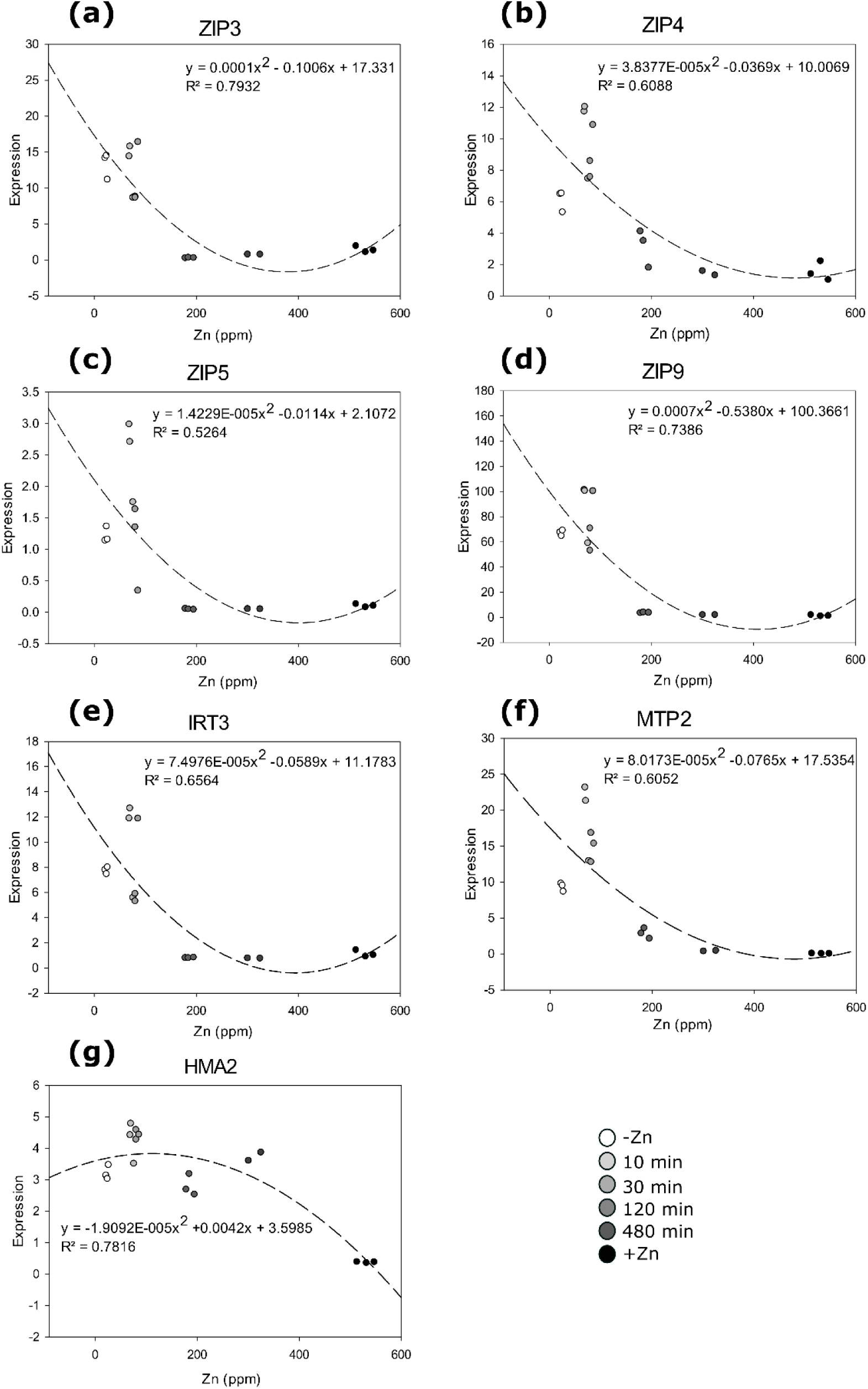
Relation between Zn levels and transcript levels of selected transporter-encoding genes in roots during the Zn starvation and re-supply time series. A quadratic equation with one unknown y= a*x*^2^ + bx + c was used for the fitting, the parameters are listed in each graph. As tissues for elemental analysis and transcript profiling were obtained in independent experiments, each biological replicate where transcript levels were measured, was plotted against a randomly selected biological replicate from the elemental analysis dataset for the same time point. Time points post re-supply are depicted in shades of grey from – Zn (white) to +Zn (black).

**Figure 8.**
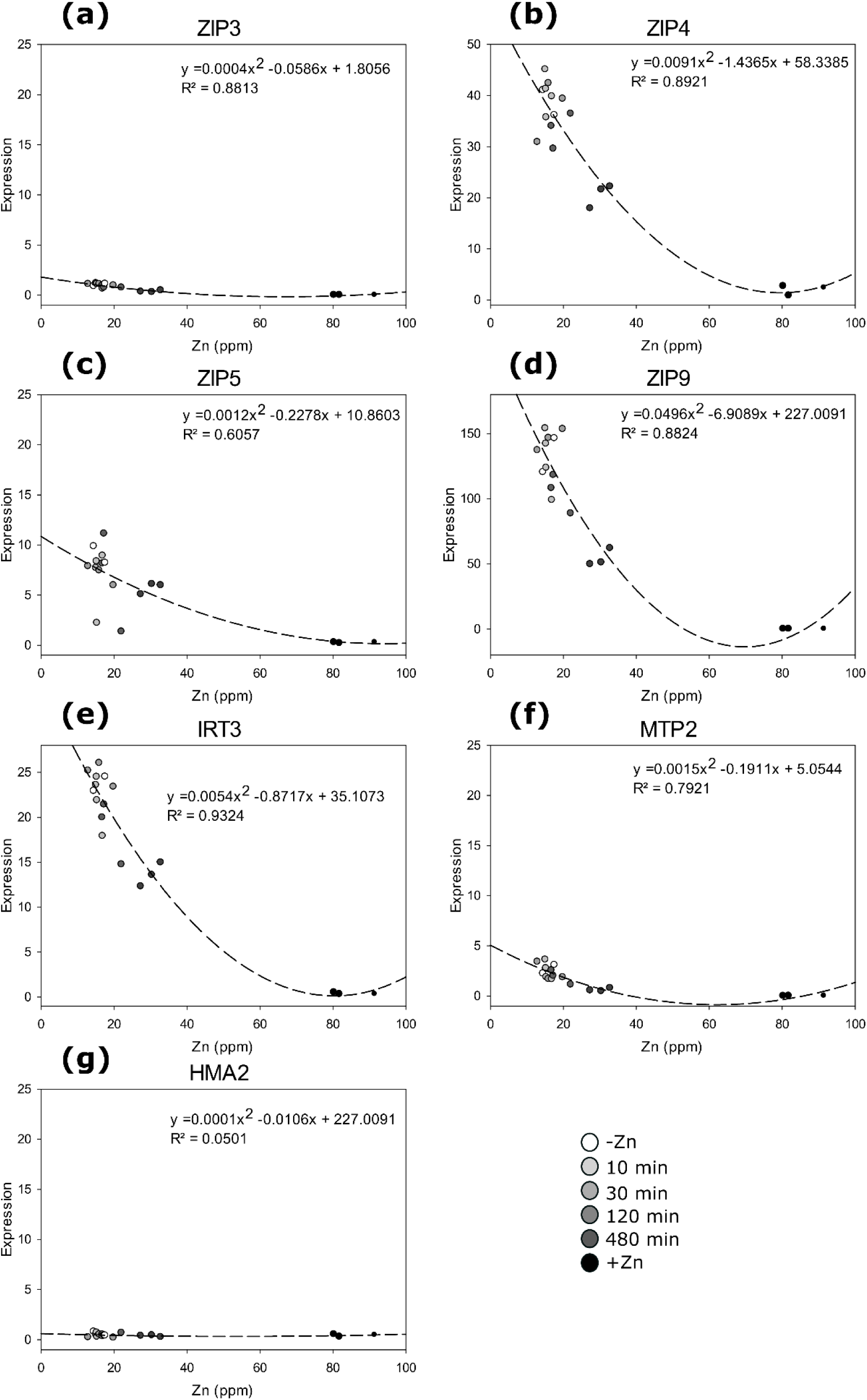
Relation between Zn levels and transcript levels of selected transporter-encoding genes in shoots during the Zn starvation and re-supply time series. A quadratic equation with one unknown y= a*x*^2^ + bx + c was used for the fitting, the parameters are listed in each graph. As tissues for elemental analysis and transcript profiling were obtained in independent experiments, each biological replicate where transcript levels were measured, was plotted against a randomly selected biological replicate from the elemental analysis dataset for the same time point. Time points post re-supply are depicted in shades of grey from – Zn (white) to +Zn (black).

The relation between Mn concentration and transcript levels was very different than for Zn (Fig. S5, S6), following a quadratic equation with an upside down parabola shape in roots. The time series was not preserved along the Mn concentration axis and, in most cases, appeared reversed. This suggested that the transient increase in transcript levels of ZIP transporters is likely responsible for the transient increase of Mn in roots at the 10 and 30min time-points. A similar pattern, although attenuated was observed in shoots (Fig. S6).

In addition to Zn, proteins primarily involved in Fe and Cu homeostasis were also dynamically regulated upon Zn deficiency and re-supply. In roots, the Ferric-chelate reductase FRO2, the iron transporter IRT1 and the vacuolar Zn transporter MTP3, all members of the FIT regulon (see Introduction, Fig. 6 i, j, k), displayed a similar expression pattern at both transcript (all 3) and protein (FRO2 and IRT1) levels, with a sinusoidal behavior along the time series (Fig. 6). This regulation pattern was similar to other ZIPs, e.g. ZIP3 or ZIP9, with the exception that FRO2, IRT1 and MTP3 were expressed at respectable levels in Zn sufficient conditions. The MTP3 protein was only detected at a few points in the time series, preventing to draw firm conclusion about its regulation, it however seemed to diverge from its regulation at transcript levels. In shoots, the transcripts of these three genes were lowly expressed, with a similar and peculiar pattern, but the corresponding proteins were not detected in agreement with their preponderant function in roots (Thomine & Vert, 2013). The YSL6 protein, a nicotianamine-metal transporter, was more highly expressed in shoots than in roots (Fig. 6l, Data S2), which is consistent with its function in Fe release from chloroplast (Divol *et al.*, 2013). However, it displayed a rapid and strong induction in roots in the early time points (10-120min.). This pattern was different from the transcript behavior, suggesting post-translational control enabling rapide response to accommodate Zn re-entry in root cells (Fig. 6l). The Vacuolar iron transporter (VIT, At4g27870), an uncharacterized transporter related to ER Mn transporters (Yamada *et al.*, 2013), showed a similar pattern (Fig. 6m). Finally, transcript and protein levels of HMA5, involved in Cu tolerance (Andres-Colas *et al.*, 2006; Kobayashi *et al.*, 2008), were higher in Zn-deficient conditions and throughout the time series compared to +Zn, with a regulation pattern very similar to HMA2 (Fig. 6n).

### Dynamics of Signaling and Regulation proteins upon Zn re-supply in roots

Coordinated metal uptake and transcriptional/translational responses of Zn transporters in roots (Fig. 7) suggested that signaling and regulation events occurred during Zn re-supply. Therefore we dissected the dynamics of proteins in the functional categories signaling, protein posttranslational modification and RNA-regulation (>4-fold change and p < 0.05, Fig.9), as putative players in cellular signal transduction pathways (Memon and Durakovic, 2014).

The signaling and regulatory response to Zn re-supply in roots progressed and amplified with time, appearing organized in two waves. First, only a few responding proteins (kinases, calcium and phosphoinositide signaling) were detected after 10min and the number of regulated proteins then moderately increased in both microsomal and soluble fractions until 120min. Second, at 480min, a massive shift of signaling processes was observed with a marked down-regulation of proteins in the microsomal fraction (103 down vs 1 up) and a reverse behavior in the soluble fraction (40 up vs 6 down) (Fig 9).

**Figure 9.**
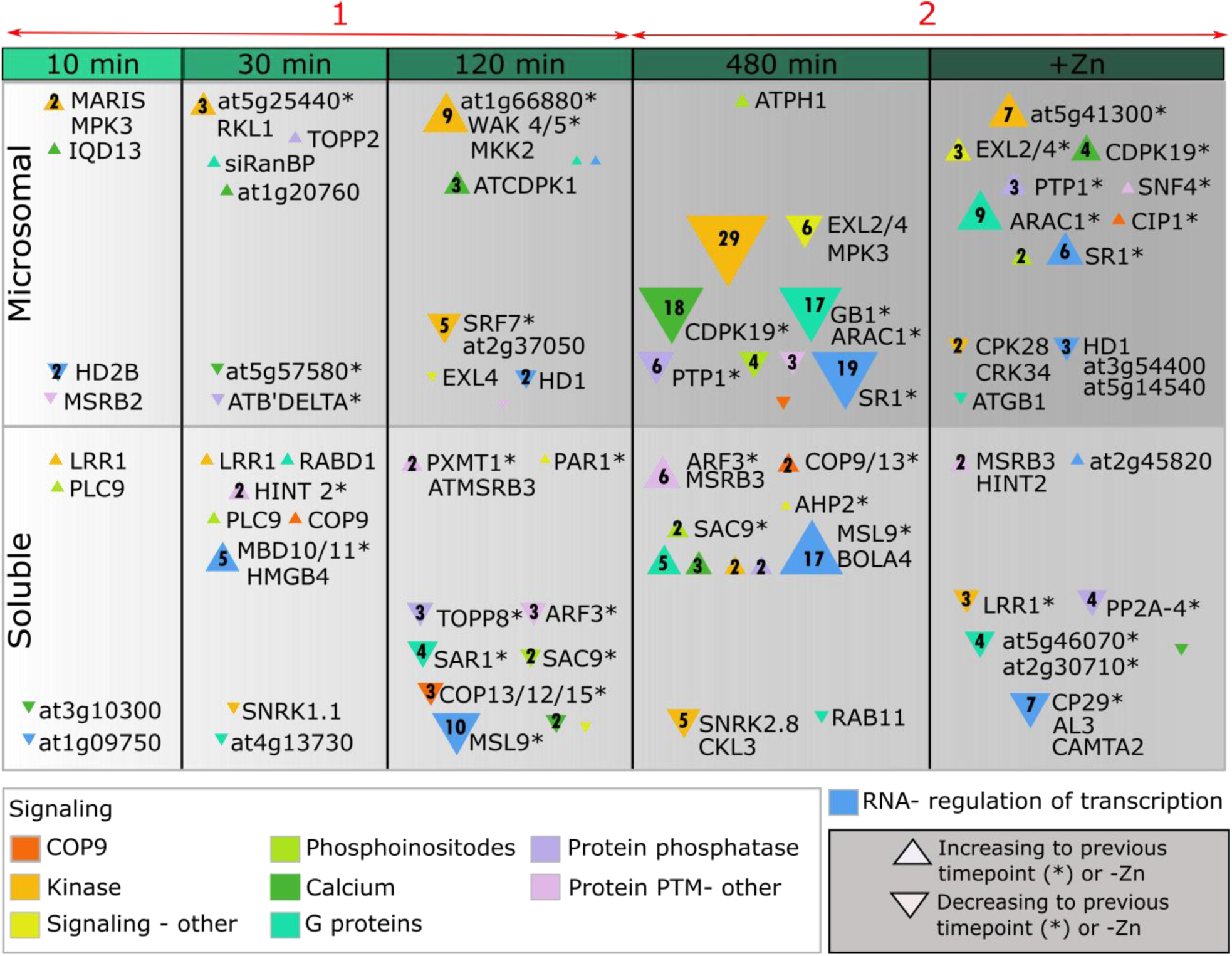
Involvement in signaling and potential regulation proteins through time during the Zn starvation and re-supply time series. Candidates were collected from the MapMan functional categories “signaling”, protein post-translational modifications and RNA regulation of transcription that passed the 4 fold change and p<0.05 thresholds (after multiple testing correction). Triangles indicate protein increase, inverted triangles protein decrease. The size of the symbols is proportional to the number of proteins from the respective functional class, and this number is also indicated in the symbol. Selected proteins, chosen from the highest responding proteins in the specific functional class, are listed by name, whenever possible, or AGI code. An asterisk (*) behind the protein name indicates that this particular protein shows >4 fold change to the previous time point. Colors indicate functional category. Orange: COP9 signalosome, yellow: kinases (mainly receptor-like kinases) and PTM signaling (MapMan categories 30 and 29.4), neon yellow: other signaling molecules related to sugar, nutrients and light (30.1 and 30.11), light green: Phosphoinositide signaling (30.4), dark green: Calcium signaling (30.3), cyan: G proteins signaling (bin 30.5), light purple: phosphatases (bin 29.4), light pink: other PTM (bin 29.4), blue: Regulation of transcription (27.3.54). The two signaling waves are indicated in red arrows. Protein names: At3g59690 (IQD13), At5g16590 (LRR1), At1g02090 (COP15), At1g07360 (COP 12), At1g18890 (ATCDPK1), At1g21210 (WAK4), At1g31160 (HINT 2), At1g48480 (RKL1), At1g56330 (SAR1), At1g66700 (PXMT1), At1g71860 (PTP1), At1g75170 (SEC14), At2g17560 (HMGB4), At2g17800 (ARAC1), At2g22300 (SR1), At2g24765 (ARF3), At2g29700 (ATPH1), At2g41970 (MARIS), At2g47640 (snRNP), At3g01090 (SNRK1.1), At3g11730 (RABD1), At3g14350 (SRF7), At3g15790 (MBD11), At3g26030 (ATB’ DELTA), At3g47220 (PLC9), At3g54040 (PAR1), At3g59770 (SAC9), At4g04800 (ATMSRB3), At4g14110 (COP9), At4g21860 (MSRB2), At4g29810 (MKK2), At4g38130 (HD1), At5g09440 (EXL4), At5g14250 (COP13), At5g16590 (LRR1), At5g19450 (CDPK19), At5g19520 (MSL9), At5g22650 (HD2B), At5g27840 (TOPP8), At5g52210 (ATGB1), At1g07140.1 (siRanBP), At5g59160 (TOPP2), At5g64260 (EXL2), At3g45640 (MPK3).

For instance, the number of responding kinases increased from 3 after 10min to more than 35 after 480min of Zn re-supply (Fig. 9). The early responding kinases included MARIS, a root hair expressed receptor-like kinase (Boisson-Dernier *et al.*, 2015), and MPK3 suggesting the inclusion of the MAP kinase pathway (Wengier *et al.*, 2018; Zhu *et al.*, 2019), which is confirmed at 120min by the detection of MKK2 (Furuya *et al.*, 2014). In addition, LRR1, a leucine rich repeat receptor kinase involved in defense signaling (Choi *et al.*, 2012) is repeatedly one of the most highly responding kinases at 10, 30min and Zn sufficiency. Interestingly, At5g67380 (AtCk2) decreased in roots at 480min (Data S6), which is homologous to the only kinase (CK2, casein kinase 2) shown to regulate the human ZIP7 protein (Taylor *et al.*, 2012).

Out of 29 responding GTP-binding proteins, 17 decreases in the microsomal fraction at 480min while 5 increased in the soluble fraction. Some of the highest responding G-proteins included siRANBP and RABD1 (30min), SAR1 (120min); GB1 and RAB11 (480min) and ARAC1 (480min and +Zn) (Fig. 9).

While individual calcium-related signaling molecules were detected earlier in the time series, the 480min time-point marked a hotspot for downregulation of Ca^2+^ signaling with 18 proteins decreasing. These included: Calcium-dependent protein kinase 19 (CDPK19), Calcium dependent protein kinase 1 (CDPK1), calmodulin-domain protein kinase 7 (CPK 7), CPK27, CPK29, CPK31, CPK32, and Calcium ATPase 2 (ACA2) (Data S6).

Among the proteins dynamically responding throughout the Zn re-supply time series, five were COP9 signalosome proteins (COP9, 12, 13, 15 and CIP1) (Wei et al., 2008). Similar to bZIP19 and bZIP23 (Fig. S2), the transcript levels of *COP9, 12, 13, 15* and *CIP1* were not significantly regulated by Zn status even though the corresponding proteins fluctuated through time (Fig. S7), suggesting that post-transcriptional regulation has a significant role in the early stages of Zn re-supply. Out of five checked proteins, four (COP9, COP12, COP15 and CIP1) showed large fluctuations in protein levels in the root soluble fraction, all having a first peak at 30min and a second at 480min. This synchronized response in roots leads to speculation that the proteasome and COP9 signalosome may have a function in establishing Zn homeostasis.

Similarly, a putative transcription factor (At1g02080) showed no significant regulation of transcript levels, but large variation in microsomal and soluble protein levels in roots and shoots (Fig S2b). Overall, a substantial number of transcription factors were regulated through the Zn re-supply time series (in blue in Fig. 9).

## Discussion

Plants and plant roots are inherently dynamic in their response to the environment with strategies ranging from molecular to anatomical changes (Arsova, et al., submitted). We postulated that this acclimation period represents the true plasticity of an organism and must be studied in detail to understand Zn uptake and homeostasis. Several studies used proteomics to clarify Zn homeostasis. Inaba et al. (2015) and (Zargar *et al.*, 2015b)Zargar et al. (2015b) are the only studies clearly focusing on Zn deficiency, whereas others (Fukao *et al.*, 2009; Fukao *et al.*, 2011; Zargar *et al.*, 2015b) focused on Zn excess or crosstalk with other metals. All studies examined steady-state protein levels at a single time point. Here, our experimental design, inspired by Talke et al. (2006) and Engelsberger and Schulze (2012), relied on consecutive sampling of treated plants to resolve sequential and dynamic changes in the Arabidopsis proteome in response to changes in Zn supply.

### A time-dependent dataset

We quantified over 6400 proteins (5249 in roots and 4698 in shoots, respectively), effectively combining 4 time-resolved datasets (Figure 2), providing one of the most encompassing proteomic studies in Arabidopsis (Majeran *et al.*, 2018). We expected that a fixed condition comparison (i.e. +Zn vs -Zn) would only reveal a fraction of the proteins responding to a change in Zn status. Indeed, with our very conservative thresholds (4-fold changes, adjusted p<0.05), only 70 proteins were regulated in the microsomal fractions between -Zn and +Zn conditions. This number increased for instance to 134 and 531 regulated proteins in -Zn/120min and -Zn/480min comparisons, respectively, depicting the multitude of dynamic processes that take place between static conditions.

Sequential sampling additionally enabled visualizing the progression of the response through the plant in context of time (Fig. 4, 5), for instance revealing points of cross-talk to other nutrients. Hence, 5 nitrogen related proteins (1 glutamate dehydrogenase, 3 glutamate synthases and nitrate reductase) and 22 proteins related to amino acid metabolism were up-regulated at 480min of Zn re-supply (Data S6). Moreover, the SnRK1.1 kinase [Sucrose non-fermenting 1 (SNF1)-related protein kinase 1.1] known to regulate both carbon and nitrogen metabolism was already up-regulated 10min post re-supply (Fig. 9) (Coello & Martinez-Barajas, 2014), highlighting a rapid impact of change in Zn supply on nitrogen homeostasis. Interestingly, the activity of SnRK2 proteins has been linked to the regulation of Fe and Cd uptake and distribution in plants (Fan *et al.*, 2014; Wang *et al.*, 2019)(see SnRK2.8, Fig. 9). Similar step by step increase in the number of responding proteins was observed for functional categories such as lipid metabolism, stress, protein or signaling (Fig. 5a).

### Time-resolution of transporter dynamics and their possible regulation

Hypothetically, a Zn-deficient plant would perform several functions upon Zn re-supply before Zn homeostasis is re-adjusted to a new, Zn-sufficient, steady-state: (i) sensing that Zn is now present, (ii) taking up Zn from the environment into roots, but also temporarily accommodating a possible Zn excess, (iii) signaling the rest of the plant that Zn is coming, (iv) eventually shutting down Zn import, (v) transporting chelated Zn from roots to shoots; (vi) distributing Zn in tissues and organic compounds/target proteins.

How Zn is sensed in the rhizosphere and in plant tissues remains an open question, with the hypothesis that the bZIP19 and bZIP23 transcription factors may partially fill this role (Assunção *et al.*, 2013). The regulation of several transporters and proteins involved in Zn and metal homeostasis upon Zn re-supply was revealed here in unprecedented detail, together with the corresponding transcript levels (Fig. 6) and the dynamics of Zn entry in the plant (Fig. 7). For instance, many ZIP transporters, already strongly up-regulated at Zn deficiency, further peaked in expression at 10-30min upon Zn re-supply (Fig. 6), before going down, reversely proportional to increased Zn concentration in roots (Fig. 7, S5). It is clear that time and/or metal amount are needed for the transporters to decrease in expression (30-120min), suggesting that a feedback mechanism inside root cells responds to either Zn ions or chelated Zn and starts a regulatory cascade once a given concentration (slightly below 200 ppm in roots) is reached. This signal may be part of a damped oscillator response (Fukuda *et al.*, 2013; Gould *et al.*, 2018) that corrects a transcription overshoot at early time points of the re-supply, and manifested as peaks of expression of many transporters at 10-30min (Fig. 6).

Sinclair et al. (2018) showed that among the genes transcriptionally regulated by Zn deficiency in Arabidopsis roots, some (e.g. *ZIPs*) are controlled by a local signal, possibly via bZIP19 and bZIP23, whereas others (e.g. *HMA2* and *MTP2*) are controlled by an unidentified shoot-born Zn deficiency signal. The dynamics of HMA2 and MTP2 transcript and protein levels upon Zn re-supply of Zn-deficient plants (Fig. 6) indicate that their regulation might be even more complex: their rapid up-then down-regulation in roots before Zn has reached shoots (Fig. 1, 6) suggests that their expression is also controlled by a local Zn sufficiency/excess signal, which overrides the systemic shoot Zn-deficiency signaling.

The observation that Zn uptake and constant accumulation is accompanied by an initial and transient (10-30min) increase in Mn concentrations in roots (Fig. 1c,e) possibly highlights the dual affinity of a number of ZIP transporters for both Zn and Mn (Milner *et al.*, 2013), but also suggests that a possible Mn excess is rapidly counteracted by exclusion. Such a Mn exclusion transporter and a possible Mn sensing mechanism remain to be identified. In contrast, Zn deficiency resulted in an increased Fe and Cu accumulation and Zn-resupply rapidly transiently decreased Fe and Cu concentrations in roots (Fig. 1), suggesting competition for uptake with Zn. Several transporters involved in Fe and Cu homeostasis were rapidly and dynamically regulated by Zn re-supply (Fig. 6h-n).

### Early signaling events upon Zn re-supply in roots

The dynamics of signaling proteins responding to Zn re-supply in roots segregated in two waves (Fig. 9, Data S6). Coincidently, the timing of the first wave matched the downregulation of Zn transporter transcript and protein levels and the increase of Zn concentration to about 200ppm (Fig. 7, 9). We speculate that information/sensing of restored Zn availability and an activation of uptake mechanisms is transmitted through some of the early responders (e.g. MARIS, MPK3, LRR1) to Zn re-supply whereas this initial response is rapidly replaced by signaling for shutting down Zn transporters, which probably involves molecules identified between 30 - 120min (Fig. 9). The second wave, which is characterized by massive downregulation of microsomal regulatory proteins and increase of soluble fraction signaling, coincided with the first increase of Zn in shoot and possibly the initial steps of restoring a Zn-sufficiency steady-state in roots. Thus, this is the time point when we expect systemic signals to become active (Sinclair *et al.*, 2018), while the roots are actively working to avoid excess Zn (Fig. 6).

Overall, this dynamic response to Zn re-supply mobilized numerous signaling proteins, including kinases, GTP-binding proteins, calcium and phosphoinositide signaling, as well as proteins of the signalosome and transcription factors (Fig. 9).

Interestingly, among the earliest responding proteins (Fig. 9), the number of identified kinases largely outnumbered phosphatases (e.g. Hint 2, TOPP2, TOPP 8, PTP1, and PP2A-4), supporting a previous claim that a variety of kinases are balanced by a smaller number of phosphatases (Smoly *et al.*, 2017). Mostly responding at 120-480min (Fig. 9), GTP-based membrane receptors activate further signaling cascades in the cytosolic fraction (Memon & Durakovic, 2014), whereas phosphoinositides are involved in multiple aspects of cellular regulation of vesicular trafficking, lipid distribution, metabolism, as well as of ion channels, pumps and transporters. For instance, the localization of NRAMP1 in Arabidopsis is controlled by the Phosphatidylinositol 3-phosphate–binding protein (Agorio *et al.*, 2017). Membrane-bound phosphoinositides are also involved in the production of the secondary messenger molecules Inositol trisphosphate (IP3) and diacylglycerol (DAG), through Phospholipase C (PLC) enzymes. The activation of PLCs in humans precedes cytoplasmic Ca signaling. G-proteins have been suggested as regulators of PLCs (Balla, 2013).

In plants, the COP9 signalosome is essential for correct expression of Fe homeostasis genes in Arabidopsis, although the corresponding COP9 genes did not respond to Fe deficiency (Eroglu & Aksoy, 2017). Several proteins of the COP9 signalosome dynamically responded to a change in Zn supply, with however often similar protein levels in Zn deficiency and sufficiency conditions (Fig. S7). The COP9 functions are associated with de-ubiquitination activity as well as protein kinase activity of various signaling regulators (Wei & Deng, 2003; Wei *et al.*, 2008). Their function could be important in understanding protein turnover of Zn- or metal-related proteins (Fig. 6), similar to the mechanism involved in IRT1 turnover (Kerkeb *et al.*, 2008; Dubeaux *et al.*, 2018).

Finally, progressive regulation of various transcription factors and proteins otherwise involved in regulation of RNA transcription culminated at 480min. It is expected that some of these will be involved in control of Zn homeostasis or of pathways which are in cross-talk to Zn homeostasis (Khan *et al.*, 2014; Naeem *et al.*, 2018).

In conclusion, our dataset identifies a number of candidates for further investigation of Zn related signaling, which will need to be confirmed by targeted mutant analyses. Admittedly, information on protein posttranslational modifications would be useful to further elucidate the rapid dynamics of metal transporters (e.g. ZIPs), similar to the human ZIP7 (Taylor *et al.*, 2012). This effort is ongoing. Directed protein interaction assays will also be necessary to fully elucidate the signal transduction cascades involved in Zn homeostasis. Overall, the time-related sampling allowed unraveling the protein dynamics upon changes in Zn supply, including cases where the start and end protein levels are similar but the protein fluctuates through time.

## Acknowledgements

Björn Usadel is thanked for constructive discussions and functional enrichment tool. B.A. and M.H. acknowledge funding as post-doctoral researcher (grant 1209413F) and research associate of F.R.S.-FNRS, respectively. Funding is provided by FNRS (grants PDR T.0206.13, MIS-F.4511.16 and CDR J.0009.17 to M.H.). S.A. is funded by a PhD grant of FZJ.

## Author contributions

MH and BA designed the research. BA, SA, MS, DB, GM, MC, BB performed experiments. BA, MH, SA analyzed the data. EDP, MW, PM helped with data interpretation. BA made the figures. BA, MH wrote the manuscript. All authors read and approved the manuscript.

## Figure Legends

### The following Supporting Information is available for this article

Figure S1. Experimental design of the Zn deficiency and re-supply.

Figure S2. Expression of the transcription factors bZIP19, bZIP23 and At1g02080.

Figure S3. Overview of the root and shoot dataset.

Figure S4. Selection process of proteins responding to Zn re-supply

Figure S5. Relation between Mn levels and transcript levels of selected transporter-encoding genes in roots during the Zn starvation and re-supply time series.

Figure S6. Relation between Mn levels and transcript levels of selected transporter-encoding genes in shoots during the Zn starvation and re-supply time series.

Figure S7. Transcript and protein levels of the COP9 signalosome members.

Table S1. Sequence of primers for quantitative RT-PCR.

Table S2. Enrichment of MapMan functional categories among the responding proteins in roots.

Table S3. Enrichment of MapMan functional categories among the responding proteins in shoots.

Data S1. Raw file names for proteomics measurements.

Data S2. Transcript expression by qRT-PCR. Data S3. Root quantified proteins.

Data S4. Shoot quantified proteins.

Data S5. Protein ratios and multiple testing correction.

Data S6. Responding proteins and enrichment of functional categories.

## References

Agorio A, Giraudat J, Bianchi MW, Marion J, Espagne C, Castaings L, Lelievre F, Curie C, Thomine S, Merlot S. 2017. Phosphatidylinositol 3-phosphate-binding protein AtPH1 controls the localization of the metal transporter NRAMP1 in Arabidopsis. Proc Natl Acad Sci U S A 114(16): E3354–E3363.

Alloway BJ. 2009. Soil factors associated with zinc deficiency in crops and humans. Environ Geochem Health 31(5): 537–548.

Andres-Colas N, Sancenon V, Rodriguez-Navarro S, Mayo S, Thiele DJ, Ecker JR, Puig S, Penarrubia L. 2006. The Arabidopsis heavy metal P-type ATPase HMA5 interacts with metallochaperones and functions in copper detoxification of roots. Plant Journal 45(2): 225–236.

Arrivault S, Senger T, Krämer U. 2006. The Arabidopsis metal tolerance protein AtMTP3 maintains metal homeostasis by mediating Zn exclusion from the shoot under Fe deficiency and Zn oversupply. Plant Journal 46(5): 861–879.

Assunção AG, Persson DP, Husted S, Schjørring JK, Alexander RD, Aarts MG. 2013. Model of how plants sense zinc deficiency. Metallomics 5(9): 1110–1116.

Assunção AG, Schat H, Aarts MG. 2010. Regulation of the adaptation to zinc deficiency in plants. Plant signaling & behavior 5(12): 1553–1555.

Balla T. 2013. Phosphoinositides: tiny lipids with giant impact on cell regulation. Physiol Rev 93(3): 1019–1137.

Barberon M, Zelazny E, Robert S, Conéjéro G, Curie C, Friml J, Vert G. 2011. Monoubiquitin-dependent endocytosis of the iron-regulated transporter 1 (IRT1) transporter controls iron uptake in plants. Proceedings of the National Academy of Sciences 108(32): E450–E458.

Bardou P, Mariette J, Escudie F, Djemiel C, Klopp C. 2014. jvenn: an interactive Venn diagram viewer. BMC bioinformatics 15(15): 293.

Becher M, Talke IN, Krall L, Krämer U. 2004. Cross-species microarray transcript profiling reveals high constitutive expression of metal homeostasis genes in shoots of the zinc hyperaccumulator Arabidopsis halleri. Plant Journal 37(2): 251–268.

Boisson-Dernier A, Franck CM, Lituiev DS, Grossniklaus U. 2015. Receptor-like cytoplasmic kinase MARIS functions downstream of CrRLK1L-dependent signaling during tip growth. Proc Natl Acad Sci U S A 112(39): 12211–12216.

Briat JF, Rouached H, Tissot N, Gaymard F, Dubos C. 2015. Integration of P, S, Fe, and Zn nutrition signals in Arabidopsis thaliana: potential involvement of PHOSPHATE STARVATION RESPONSE 1 (PHR1). Frontiers in plant science 6: 290.

Broadley MR, White PJ, Hammond JP, Zelko I, Lux A. 2007. Zinc in plants. The New phytologist 173(4): 677–702.

Brumbarova T, Bauer P, Ivanov R. 2015. Molecular mechanisms governing Arabidopsis iron uptake. Trends in Plant Science 20(2): 124–133.

Chiapello M, Martino E, Perotto S. 2015. Common and metal-specific proteomic responses to cadmium and zinc in the metal tolerant ericoid mycorrhizal fungus Oidiodendron maius Zn. Metallomics 7(5): 805–815.

Chmielowska-Bak J, Gzyl J, Rucińska-Sobkowiak R, Arasimowicz-Jelonek M, Deckert J. 2014. The new insights into cadmium sensing. Frontiers in plant science 5: 245.

Choi DS, Hwang IS, Hwang BK. 2012. Requirement of the cytosolic interaction between PATHOGENESIS-RELATED PROTEIN10 and LEUCINE-RICH REPEAT PROTEIN1 for cell death and defense signaling in pepper. Plant Cell 24(4): 1675–1690.

Claus J, Bohmann A, Chavarria-Krauser A. 2013. Zinc uptake and radial transport in roots of Arabidopsis thaliana: a modelling approach to understand accumulation. Annals of botany 112(2): 369–380.

Coello P, Martinez-Barajas E. 2014. The activity of SnRK1 is increased in Phaseolus vulgaris seeds in response to a reduced nutrient supply. Frontiers in plant science 5: 196.

Colangelo EP, Guerinot ML. 2004. The essential basic helix-loop-helix protein FIT1 is required for the iron deficiency response. Plant Cell 16(12): 3400–3412.

Cox J, Mann M. 2008. MaxQuant enables high peptide identification rates, individualized p.p.b.-range mass accuracies and proteome-wide protein quantification. Nat Biotechnol 26(12): 1367–1372.

Deinlein U, Weber M, Schmidt H, Rensch S, Trampczynska A, Hansen TH, Husted S, Schjoerring JK, Talke IN, Kramer U, et al. 2012. Elevated Nicotianamine Levels in Arabidopsis halleri Roots Play a Key Role in Zinc Hyperaccumulation. Plant Cell 24(2): 708–723.

Desbrosses-Fonrouge A-G, Voigt K, Schröder A, Arrivault S, Thomine S, Krämer U. 2005. Arabidopsis thaliana MTP1 is a Zn transporter in the vacuolar membrane which mediates Zn detoxification and drives leaf Zn accumulation. Febs Letters 579(19): 4165–4174.

Deutsch EW, Csordas A, Sun Z, Jarnuczak A, Perez-Riverol Y, Ternent T, Campbell DS, Bernal-Llinares M, Okuda S, Kawano S, et al. 2017. The ProteomeXchange consortium in 2017: supporting the cultural change in proteomics public data deposition. Nucleic Acids Res 45(D1): D1100–D1106.

Divol F, Couch D, Conejero G, Roschzttardtz H, Mari S, Curie C. 2013. The Arabidopsis YELLOW STRIPE LIKE4 and 6 Transporters Control Iron Release from the Chloroplast. Plant Cell 25(3): 1040–1055.

Dubeaux G, Neveu J, Zelazny E, Vert G. 2018. Metal Sensing by the IRT1 Transporter-Receptor Orchestrates Its Own Degradation and Plant Metal Nutrition. Molecular Cell 69(6): 953–964 e955.

Engelsberger WR, Schulze WX. 2012. Nitrate and ammonium lead to distinct global dynamic phosphorylation patterns when resupplied to nitrogen-starved Arabidopsis seedlings. Plant Journal 69(6): 978–995.

Eroglu S, Aksoy E. 2017. Genome-wide analysis of gene expression profiling revealed that COP9 signalosome is essential for correct expression of Fe homeostasis genes in Arabidopsis. Biometals: an international journal on the role of metal ions in biology, biochemistry, and medicine 30(5): 685–698.

Fan SK, Fang XZ, Guan MY, Ye YQ, Lin XY, Du ST, Jin CW. 2014. Exogenous abscisic acid application decreases cadmium accumulation in Arabidopsis plants, which is associated with the inhibition of IRT1-mediated cadmium uptake. Frontiers in plant science 5: 721.

Farinati S, DalCorso G, Bona E, Corbella M, Lampis S, Cecconi D, Polati R, Berta G, Vallini G, Furini A. 2009. Proteomic analysis of Arabidopsis halleri shoots in response to the heavy metals cadmium and zinc and rhizosphere microorganisms. Proteomics 9(21): 4837–4850.

Fukao Y, Ferjani A, Fujiwara M, Nishimori Y, Ohtsu I. 2009. Identification of zinc-responsive proteins in the roots of Arabidopsis thaliana using a highly improved method of two-dimensional electrophoresis. Plant Cell Physiol 50(12): 2234–2239.

Fukao Y, Ferjani A, Tomioka R, Nagasaki N, Kurata R, Nishimori Y, Fujiwara M, Maeshima M. 2011. iTRAQ analysis reveals mechanisms of growth defects due to excess zinc in Arabidopsis. Plant Physiol 155(4): 1893–1907.

Fukuda H, Murase H, Tokuda IT. 2013. Controlling circadian rhythms by dark-pulse perturbations in Arabidopsis thaliana. Scientific reports 3: 1533.

Furuya T, Matsuoka D, Nanmori T. 2014. Membrane rigidification functions upstream of the MEKK1-MKK2-MPK4 cascade during cold acclimation in Arabidopsis thaliana. FEBS Lett 588(11): 2025–2030.

Gould PD, Domijan M, Greenwood M, Tokuda IT, Rees H, Kozma-Bognar L, Hall AJ, Locke JC. 2018. Coordination of robust single cell rhythms in the Arabidopsis circadian clock via spatial waves of gene expression. Elife 7.

Gravot A, Lieutaud A, Verret F, Auroy P, Vavasseur A, Richaud P. 2004. AtHMA3, a plant P-1B-ATPase, functions as a Cd/Pb transporter in yeast. Febs Letters 561(1-3): 22–28.

Hanikenne M, Baurain D. 2014. Origin and evolution of metal P-type ATPases in Plantae (Archaeplastida). Frontiers in plant science 4.

Hanikenne M, Nouet C. 2011. Metal hyperaccumulation and hypertolerance: a model for plant evolutionary genomics. Curr Opin Plant Biol 14(3): 252–259.

Hanikenne M, Talke IN, Haydon MJ, Lanz C, Nolte A, Motte P, Kroymann J, Weigel D, Krämer U. 2008. Evolution of metal hyperaccumulation required cis-regulatory changes and triplication of HMA4. Nature 453(7193): 391–395.

Huang XY, Chao DY, Gao JP, Zhu MZ, Shi M, Lin HX. 2009. A previously unknown zinc finger protein, DST, regulates drought and salt tolerance in rice via stomatal aperture control. Genes & development 23(15): 1805–1817.

Hussain D, Haydon MJ, Wang Y, Wong E, Sherson SM, Young J, Camakaris J, Harper JF, Cobbett CS. 2004. P-type ATPase heavy metal transporters with roles in essential zinc homeostasis in Arabidopsis. The Plant Cell 16(5): 1327–1339.

Inaba S, Kurata R, Kobayashi M, Yamagishi Y, Mori I, Ogata Y, Fukao Y. 2015. Identification of putative target genes of bZIP19, a transcription factor essential for Arabidopsis adaptation to Zn deficiency in roots. Plant J 84(2): 323–334.

Kerkeb L, Mukherjee I, Chatterjee I, Lahner B, Salt DE, Connolly EL. 2008. Iron-induced turnover of the Arabidopsis IRON-REGULATED TRANSPORTER1 metal transporter requires lysine residues. Plant Physiol 146(4): 1964–1973.

Khan GA, Bouraine S, Wege S, Li Y, de Carbonnel M, Berthomieu P, Poirier Y, Rouached H. 2014. Coordination between zinc and phosphate homeostasis involves the transcription factor PHR1, the phosphate exporter PHO1, and its homologue PHO1;H3 in Arabidopsis. Journal of Experimental Botany 65(3): 871–884.

Kierszniowska S, Walther D, Schulze WX. 2009. Ratio-dependent significance thresholds in reciprocal 15N- labeling experiments as a robust tool in detection of candidate proteins responding to biological treatment. Proteomics 9(7): 1916–1924.

Kobayashi Y, Kuroda K, Kimura K, Southron-Francis JL, Furuzawa A, Kimura K, Iuchi S, Kobayashi M, Taylor GJ, Koyama H. 2008. Amino acid polymorphisms in strictly conserved domains of a P-type ATPase HMA5 are involved in the mechanism of copper tolerance variation in Arabidopsis. Plant Physiology 148(2): 969–980.

Kodaira KS, Qin F, Tran LS, Maruyama K, Kidokoro S, Fujita Y, Shinozaki K, Yamaguchi-Shinozaki K. 2011. Arabidopsis Cys2/His2 zinc-finger proteins AZF1 and AZF2 negatively regulate abscisic acidrepressive and auxin-inducible genes under abiotic stress conditions. Plant Physiol 157(2): 742–756.

Krämer U, Talke IN, Hanikenne M. 2007. Transition metal transport. FEBS Lett 581(12): 2263–2272.

Lucini L, Bernardo L. 2015. Comparison of proteome response to saline and zinc stress in lettuce. Frontiers in plant science 6: 240.

Luo Z-B, He J, Polle A, Rennenberg H. 2016. Heavy metal accumulation and signal transduction in herbaceous and woody plants: paving the way for enhancing phytoremediation efficiency. Biotechnol Adv 34(6): 1131–1148.

Majeran W, Le Caer JP, Ponnala L, Meinnel T, Giglione C. 2018. Targeted Profiling of Arabidopsis thaliana Subproteomes Illuminates Co- and Posttranslationally N-Terminal Myristoylated Proteins. Plant Cell 30(3): 543–562.

Masood A, Iqbal N, Khan NA. 2012. Role of ethylene in alleviation of cadmium-induced photosynthetic capacity inhibition by sulphur in mustard. Plant, Cell & Environment 35(3): 524–533.

Memon AR, Durakovic C. 2014. Signal Perception and Transduction in Plants. Periodicals of Engineering and Natural Sciences (PEN) 2(2): 15–29.

Milner MJ, Seamon J, Craft E, Kochian LV. 2013. Transport properties of members of the ZIP family in plants and their role in Zn and Mn homeostasis. Journal of Experimental Botany 64(1): 369–381.

Morel M, Crouzet J, Gravot A, Auroy P, Leonhardt N, Vavasseur A, Richaud P. 2009. AtHMA3, a P-1B- ATPase Allowing Cd/Zn/Co/Pb Vacuolar Storage in Arabidopsis. Plant Physiology 149(2): 894–904.

Murashige T, Skoog F. 1962. A Revised Medium for Rapid Growth and Bio Assays with Tobacco Tissue Cultures. Physiologia Plantarum 15(3): 473–497.

Naeem A, Aslam M, Lodhi A. 2018. Improved potassium nutrition retrieves phosphorus-induced decrease in zinc uptake and grain zinc concentration of wheat. J Sci Food Agric 98(11): 4351–4356.

Nouet C, Charlier JB, Carnol M, Bosman B, Farnir F, Motte P, Hanikenne M. 2015. Functional analysis of the three HMA4 copies of the metal hyperaccumulator Arabidopsis halleri. Journal of Experimental Botany.

Nouet C, Motte P, Hanikenne M. 2011. Chloroplastic and mitochondrial metal homeostasis. Trends in Plant Science 16(7): 395–404.

Palmer CM, Guerinot ML. 2009. Facing the challenges of Cu, Fe and Zn homeostasis in plants. Nature Chemical Biology 5(5): 333–340.

Palmgren MG, Clemens S, Williams LE, Krämer U, Borg S, Schjorring JK, Sanders D. 2008. Zinc biofortification of cereals: problems and solutions. Trends in Plant Science 13(9): 464–473.

Perez-Riverol Y, Csordas A, Bai J, Bernal-Llinares M, Hewapathirana S, Kundu DJ, Inuganti A, Griss J, Mayer G, Eisenacher M, et al. 2019. The PRIDE database and related tools and resources in 2019: improving support for quantification data. Nucleic Acids Res 47(D1): D442–D450.

Pfaffl MW, Lange IG, Daxenberger A, Meyer HHD. 2001. Tissue-specific expression pattern of estrogen receptors (ER): Quantification of ER alpha and ER beta mRNA with real-time RT-PCR. Apmis 109(5): 345–355.

Ricachenevsky FK, Menguer PK, Sperotto RA, Fett JP. 2015. Got to hide your Zn away: Molecular control of Zn accumulation and biotechnological applications. Plant Sci 236: 1–17.

Schwacke R, Ponce-Soto GY, Krause K, Bolger AM, Arsova B, Hallab A, Gruden K, Stitt M, Bolger ME, Usadel B. 2019. MapMan4: a refined protein classification and annotation framework applicable to multi-omics data analysis. Molecular Plant.

Sharma SS, Dietz KJ, Mimura T. 2016. Vacuolar compartmentalization as indispensable component of heavy metal detoxification in plants. Plant Cell and Environment 39(5): 1112–1126.

Sinclair SA, Krämer U. 2012. The zinc homeostasis network of land plants. Biochim Biophys Acta 1823(9): 1553–1567.

Sinclair SA, Senger T, Talke IN, Cobbett CS, Haydon MJ, Kraemer U. 2018. Systemic upregulation of MTP2- and HMA2-mediated Zn partitioning to the shoot supplements local Zn deficiency responses of Arabidopsis. Plant Cell.

Smoly I, Shemesh N, Ziv-Ukelson M, Ben-Zvi A, Yeger-Lotem E. 2017. An Asymmetrically Balanced Organization of Kinases versus Phosphatases across Eukaryotes Determines Their Distinct Impacts. PLoS Comput Biol 13(1): e1005221.

Talke IN, Hanikenne M, Krämer U. 2006. Zinc-dependent global transcriptional control, transcriptional deregulation, and higher gene copy number for genes in metal homeostasis of the hyperaccumulator Arabidopsis halleri. Plant Physiol 142(1): 148–167.

Taylor KM, Hiscox S, Nicholson RI, Hogstrand C, Kille P. 2012. Protein kinase CK2 triggers cytosolic zinc signaling pathways by phosphorylation of zinc channel ZIP7. Science signaling 5(210): ra11.

Thomine S, Vert G. 2013. Iron transport in plants: better be safe than sorry. Curr Opin Plant Biol 16(3): 322–327.

Usadel B, Poree F, Nagel A, Lohse M, Czedik-Eysenberg A, Stitt M. 2009. A guide to using MapMan to visualize and compare Omics data in plants: a case study in the crop species, Maize. Plant Cell and Environment 32(9): 1211–1229.

van de Mortel JE, Almar Villanueva L, Schat H, Kwekkeboom J, Coughlan S, Moerland PD, Ver Loren van Themaat E, Koornneef M, Aarts MG. 2006. Large expression differences in genes for iron and zinc homeostasis, stress response, and lignin biosynthesis distinguish roots of Arabidopsis thaliana and the related metal hyperaccumulator Thlaspi caerulescens. Plant Physiol 142(3): 1127–1147.

Vert G, Barberon M, Zelazny E, Séguéla M, Briat J-F, Curie C. 2009. Arabidopsis IRT2 cooperates with the high-affinity iron uptake system to maintain iron homeostasis in root epidermal cells. Planta 229(6): 1171–1179.

Vert G, Grotz N, Dédaldéchamp F, Gaymard F, Guerinot ML, Briat J-F, Curie C. 2002. IRT1, an Arabidopsis transporter essential for iron uptake from the soil and for plant growth. The Plant Cell 14(6): 1223–1233.

Wang R, Wang C, Yao Q, Xiao X, Fan X, Sha L, Zeng J, Kang H, Zhang H, Zhou Y, et al. 2019. The polish wheat (Triticum polonicum L.) TpSnRK2.10 and TpSnRK2.11 meditate the accumulation and the distribution of cd and Fe in transgenic Arabidopsis plants. BMC genomics 20(1): 210.

Wang R, Wang J, Zhao L, Yang S, Song Y. 2015. Impact of heavy metal stresses on the growth and auxin homeostasis of Arabidopsis seedlings. Biometals: an international journal on the role of metal ions in biology, biochemistry, and medicine 28(1): 123–132.

Wei N, Deng XW. 2003. The COP9 signalosome. Annu Rev Cell Dev Biol 19: 261–286.

Wei N, Serino G, Deng XW. 2008. The COP9 signalosome: more than a protease. Trends Biochem Sci 33(12): 592–600.

Wengier DL, Lampard GR, Bergmann DC. 2018. Dissection of MAPK signaling specificity through protein engineering in a developmental context. BMC Plant Biol 18(1): 60.

Williams LE, Mills RF. 2005. P-1B-ATPases - an ancient family of transition metal pumps with diverse functions in plants. Trends in Plant Science 10(10): 491–502.

Wintz H, Fox T, Wu YY, Feng V, Chen W, Chang HS, Zhu T, Vulpe C. 2003a. Expression profiles of Arabidopsis thaliana in mineral deficiencies reveal novel transporters involved in metal homeostasis. The Journal of Biological Chemistry 278(48): 47644–47653.

Wintz H, Fox T, Wu YY, Feng V, Chen W, Chang HS, Zhu T, Vulpe C. 2003b. Expression profiles of Arabidopsis thaliana in mineral deficiencies reveal novel transporters involved in metal homeostasis. J Biol Chem 278(48): 47644–47653.

Yamada K, Nagano AJ, Nishina M, Hara-Nishimura I, Nishimura M. 2013. Identification of two novel endoplasmic reticulum body-specific integral membrane proteins. Plant Physiol 161(1): 108–120.

Yan JY, Li CX, Sun L, Ren JY, Li GX, Ding ZJ, Zheng SJ. 2016. A WRKY Transcription Factor Regulates Fe Translocation under Fe Deficiency. Plant Physiology 171(3): 2017–2027.

Zargar SM, Fujiwara M, Inaba S, Kobayashi M, Kurata R, Ogata Y, Fukao Y. 2015a. Correlation analysis of proteins responsive to Zn, Mn, or Fe deficiency in Arabidopsis roots based on iTRAQ analysis. Plant Cell Reports 34(1): 157–166.

Zargar SM, Kurata R, Inaba S, Oikawa A, Fukui R, Ogata Y, Agrawal GK, Rakwal R, Fukao Y. 2015b. Quantitative proteomics of Arabidopsis shoot microsomal proteins reveals a cross-talk between excess zinc and iron deficiency. Proteomics 15(7): 1196–1201.

Zauber H, Schulze WX. 2012. Proteomics wants cRacker: automated standardized data analysis of LC-MS derived proteomic data. J Proteome Res 11(11): 5548–5555.

Zhu Q, Shao Y, Ge S, Zhang M, Zhang T, Hu X, Liu Y, Walker J, Zhang S, Xu J. 2019. A MAPK cascade downstream of IDA-HAE/HSL2 ligand-receptor pair in lateral root emergence. Nat Plants.

