## Supplementary material for "Resolution of the proteome, transcript and ionome dynamics upon Zn re-supply in Zn-deficient Arabidopsis": Fig. S

*New Phytologist* **Supporting <u>figures and tables</u>**

The following Supporting Information is available for this article:

Figure S1. Experimental design of the Zn deficiency and re-supply.
Figure S2. Expression of the transcription factors bZIP19, bZIP23 and At1g02080.
Figure S3. Overview of the root and shoot dataset.
Figure S4. Selection process of proteins responding to Zn re-supply
Figure S5. Relation between Mn levels and transcript levels of selected transporter-encoding genes in roots during the Zn starvation and re-supply time series.
Figure S6. Relation between Mn levels and transcript levels of selected transporter-encoding genes in shoots during the Zn starvation and re-supply time series.
Figure S7. Transcript and protein levels of the COP9 signalosome members.
Table S1. Sequence of primers for quantitative RT-PCR.
Table S2. Enrichment of MapMan functional categories among the responding proteins in roots.
Table S3. Enrichment of MapMan functional categories among the responding proteins in shoots.
Data S1. Raw file names for proteomics measurements.
Data S2. Transcript expression by qRT-PCR.
Data S3. Root quantified proteins.
Data S4. Shoot quantified proteins.
Data S5. Protein ratios and multiple testing correction.
Data S6. Responding proteins and enrichment of functional categories.

Supplemental figures:

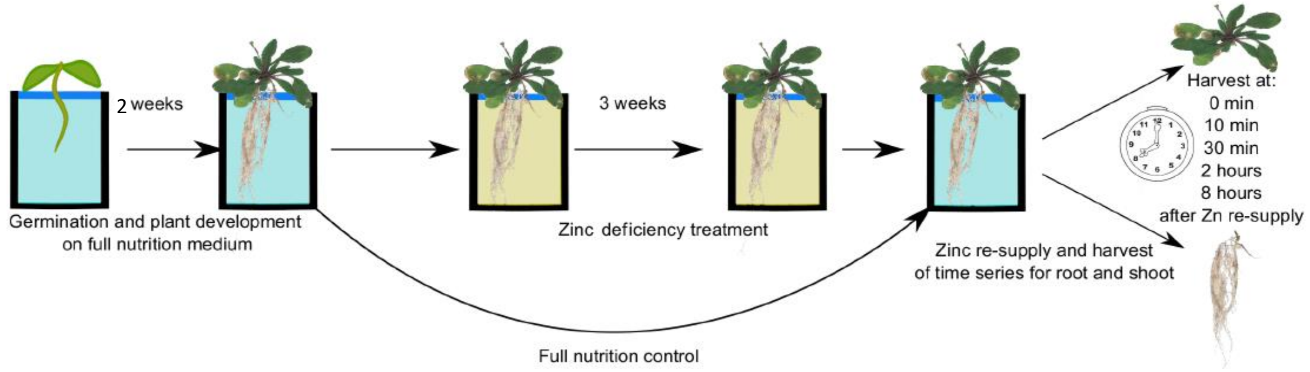


Figure S1. Experimental design of the Zn deficiency and re-supply. After germination on plates, plants were transferred in hydroponic medium with 1µM Zn for a period of 2 weeks after which they were grown for 3 weeks without Zn. After re-supply with nutrient solution containing 1µM Zn, root and shoot tissues were harvested at the indicated time-points. Zn deficiency (-Zn, 0min) and full nutrition controls were also included in the design.

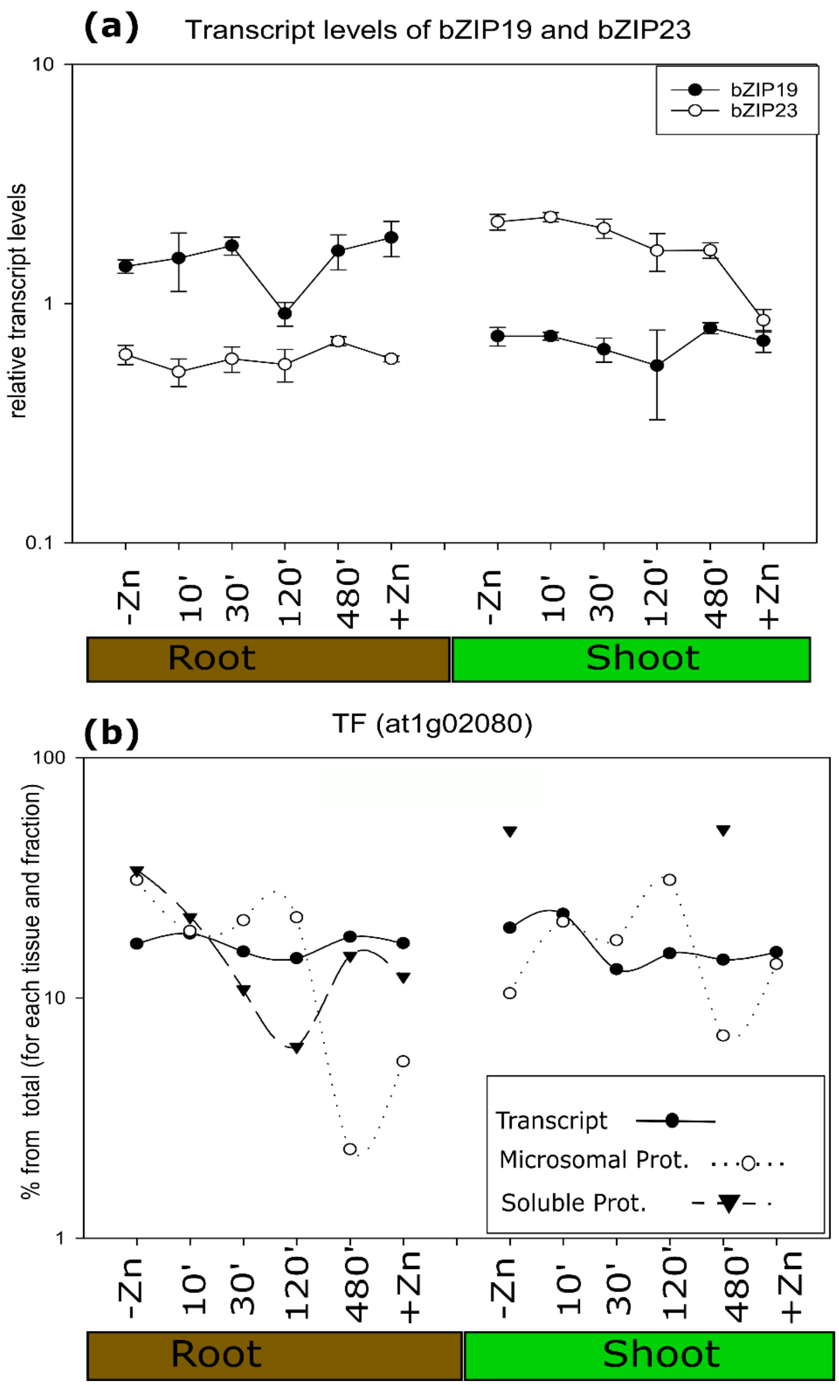


Figure S2. Expression of the transcription factors bZIP19, bZIP23 and At1g02080. Relative transcript levels of bZIP19 and bZIP23, the values are presented as 2-ddCT normalized on 2 housekeeping genes as in (Nouet et al; 2015), the value on the y-axis is common log (a). Transcript and protein levels of At1g02080. To present both transcript and protein levels for a gene on the same scale, the transcript expression/protein intensity is presented as the percent for the respective time point from the total of all time points in that tissue and fraction. The reader is reminded that these are relative values to start with and that transcript and protein intensities from each protein fraction are obtained in separate extraction steps (b).

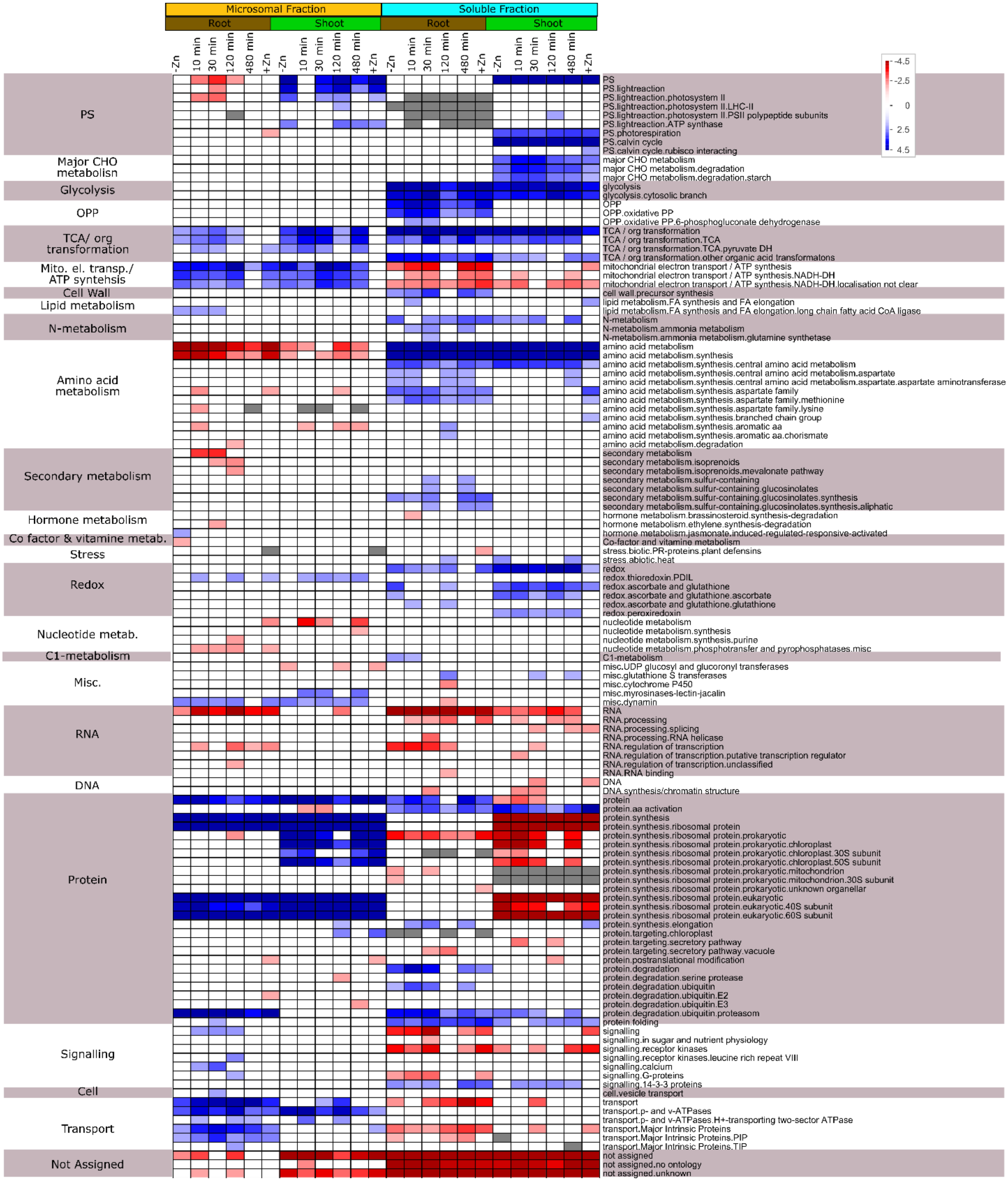

Microsomal Fraction
Soluble Fraction
Root
Shoot
Root
Shoot
-Zn
10 min
30 min
120 min
480 min
+Zn
-4.5
-2.5
0
2.5
4.5
PS
Major CHO metabolisn
Glycolysis
OPP
TCA/ org transformation
Mito. el. transp./ ATP syntehsis
Cell Wall
Lipid metabolism
N-metabolism
Amino acid metabolism
Secondary metabolism
Hormone metabolism
Co factor & vitamine metab.
Stress
Redox
Nucleotide metab.
C1-metabolism
Misc.
RNA
DNA
Protein
Signalling
Cell
Transport
Not Assigned
PS
PS.lightreaction
PS.lightreaction.photosystem II
PS.lightreaction.photosystem II.LHC-II
PS.lightreaction.photosystem II.PSII polypeptide subunits
PS.lightreaction.ATP synthase
PS.photorespiration
PS.calvin cycle
PS.calvin cycle.rubisco interacting
major CHO metabolism
major CHO metabolism.degradation
major CHO metabolism.degradation.starch
glycolysis
glycolysis.cytosolic branch
OPP
OPP.oxidative PP
OPP.oxidative PP.6-phosphogluconate dehydrogenase
TCA / org transformation
TCA / org transformation.TCA
TCA / org transformation.TCA.pyruvate DH
TCA / org transformation.other organic acid transformatons
mitochondrial electron transport / ATP synthesis
mitochondrial electron transport / ATP synthesis.NADH-DH
mitochondrial electron transport / ATP synthesis.NADH-DH.localisation not clear
cell wall.precursor synthesis
lipid metabolism.FA synthesis and FA elongation
lipid metabolism.FA synthesis and FA elongation.long chain fatty acid CoA ligase
N-metabolism
N-metabolism.ammonia metabolism
N-metabolism.ammonia metabolism.glutamine synthetase
amino acid metabolism
amino acid metabolism.synthesis
amino acid metabolism.synthesis.central amino acid metabolism
amino acid metabolism.synthesis.central amino acid metabolism.aspartate
amino acid metabolism.synthesis.central amino acid metabolism.aspartate.aspartate aminotransferase
amino acid metabolism.synthesis.aspartate family
amino acid metabolism.synthesis.aspartate family.methionine
amino acid metabolism.synthesis.aspartate family.lysine
amino acid metabolism.synthesis.branched chain group
amino acid metabolism.synthesis.aromatic aa
amino acid metabolism.synthesis.aromatic aa.chorismate
amino acid metabolism.degradation
secondary metabolism
secondary metabolism.isoprenoids
secondary metabolism.isoprenoids.mevalonate pathway
secondary metabolism.sulfur-containing
secondary metabolism.sulfur-containing.glucosinolates
secondary metabolism.sulfur-containing.glucosinolates.synthesis
secondary metabolism.sulfur-containing.glucosinolates.synthesis.aliphatic
hormone metabolism.brassinosteroid.synthesis-degradation
hormone metabolism.ethylene.synthesis-degradation
hormone metabolism.jasmonate.induced-regulated-responsive-activated
Co-factor and vitamine metabolism
stress.biotic.PR-proteins.plant defensins
stress.abiotic.heat
redox
redox.thioredoxin.PDIL
redox.ascorbate and glutathione
redox.ascorbate and glutathione.ascorbate
redox.ascorbate and glutathione.glutathione
redox.peroxiredoxin
nucleotide metabolism
nucleotide metabolism.synthesis
nucleotide metabolism.synthesis.purine
nucleotide metabolism.phosphotransfer and pyrophosphatases.misc
C1-metabolism
misc.UDP glucosyl and glucoronyl transferases
misc.glutathione S transferases
misc.cytochrome P450
misc.myrosinases-lectin-jacalin
misc.dynamin
RNA
RNA.processing
RNA.processing.splicing
RNA.processing.RNA helicase
RNA.regulation of transcription
RNA.regulation of transcription.putative transcription regulator
RNA.regulation of transcription.unclassified
RNA.RNA binding
DNA
DNA.synthesis/chromatin structure
protein
protein.aa activation
protein.synthesis
protein.synthesis.ribosomal protein
protein.synthesis.ribosomal protein.prokaryotic
protein.synthesis.ribosomal protein.prokaryotic.chloroplast
protein.synthesis.ribosomal protein.prokaryotic.chloroplast.30S subunit
protein.synthesis.ribosomal protein.prokaryotic.chloroplast.50S subunit
protein.synthesis.ribosomal protein.prokaryotic.mitochondrion
protein.synthesis.ribosomal protein.prokaryotic.mitochondrion.30S subunit
protein.synthesis.ribosomal protein.prokaryotic.unknown organellar
protein.synthesis.ribosomal protein.eukaryotic
protein.synthesis.ribosomal protein.eukaryotic.40S subunit
protein.synthesis.ribosomal protein.eukaryotic.60S subunit
protein.synthesis.elongation
protein.targeting.chloroplast
protein.targeting.secretory pathway
protein.targeting.secretory pathway.vacuole
protein.postranslational modification
protein.degradation
protein.degradation.serine protease
protein.degradation.ubiquitin
protein.degradation.ubiquitin.E2
protein.degradation.ubiquitin.E3
protein.degradation.ubiquitin.proteasom
protein.folding
signalling
signalling.in sugar and nutrient physiology
signalling.receptor kinases
signalling.receptor kinases.leucine rich repeat VIII
signalling.calcium
signalling.G-proteins
signalling.14-3-3 proteins
cell.vesicle transport
transport
transport.p- and v-ATPases
transport.p- and v-ATPases.H+-transporting two-sector ATPase
transport.Major Intrinsic Proteins
transport.Major Intrinsic Proteins.PIP
transport.Major Intrinsic Proteins.TIP
not assigned
not assigned.no ontology
not assigned.unknown

Figure S3: Overview of the root and shoot dataset. The protein levels were log 2 transformed and sorted into MapMan bins (Usadel et al., 2009), the proteins above a cut-off of (+/-) 1 were subjected to a bin-wise Wilcoxon test and the Benjamini-Hochhberg multiple testing correction. Data is presented using PageMan (Usadel et al., 2006), over represented categories (blue); under-represented categories (red)

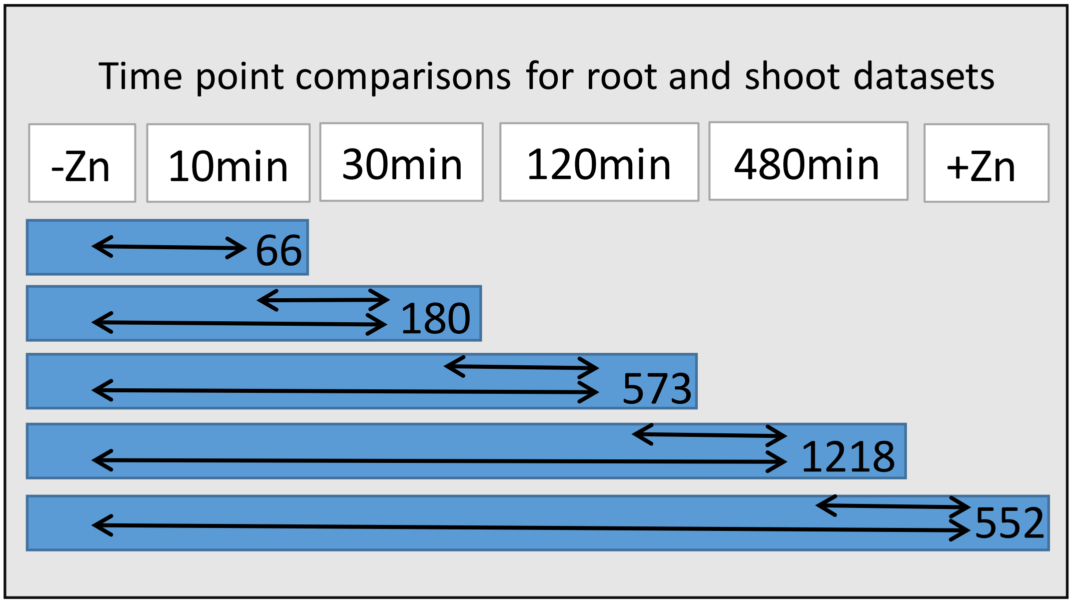


Figure S4. Selection process of proteins responding to Zn re-supply in roots. After protein quantification using cRacker and calculation of fold changes between (i) consecutive time-points and (ii) Zn deficiency and each time point (Zauber and Schulze 2012), proteins with a minimum of 4-fold change in either of these comparisons (and adjusted $p<0.05$) were merged from both fractions to tissue level in Data S6. The numbers in the blue boxes represent the root specific proteins.

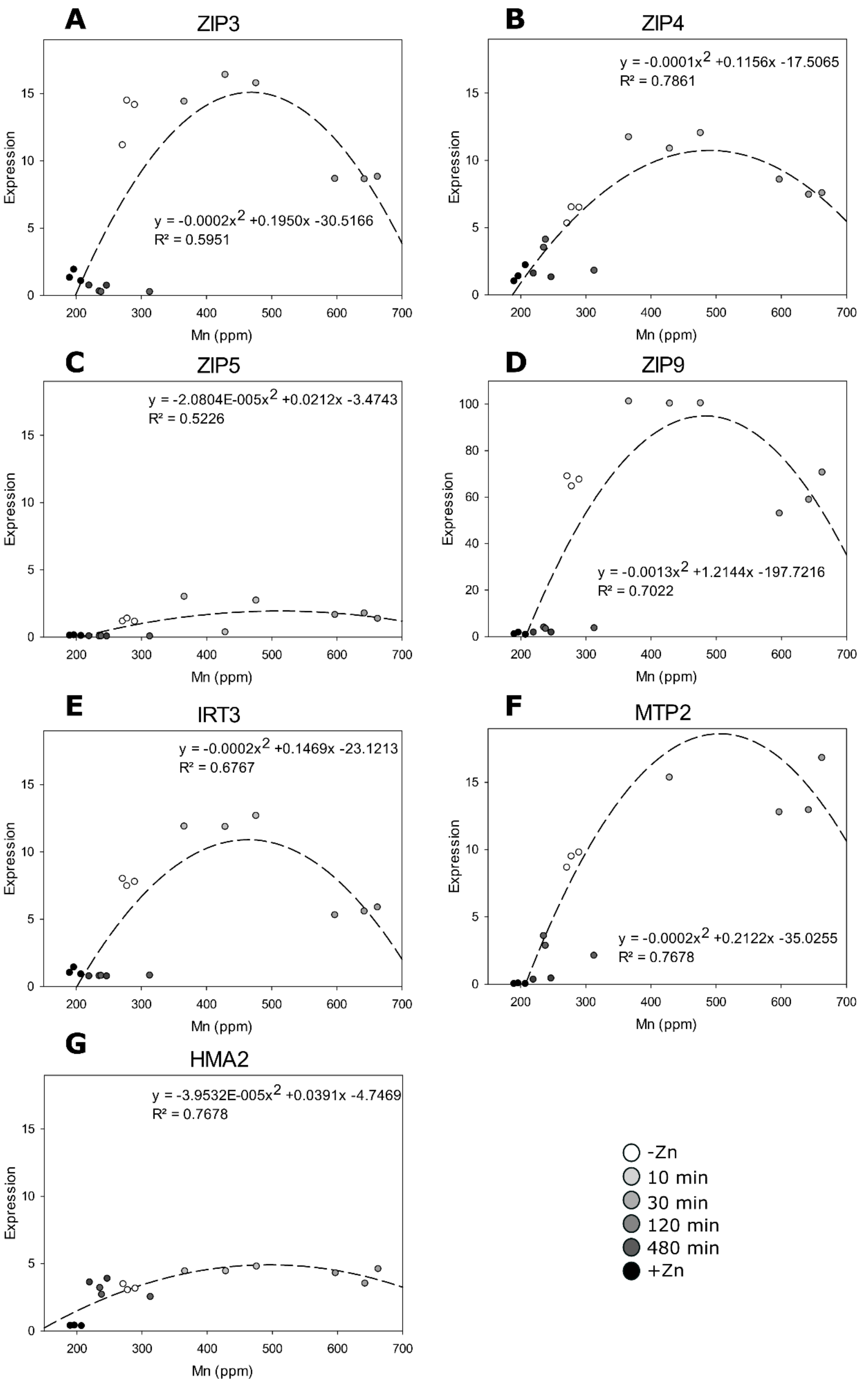


Figure S5. Relation between Mn levels and transcript levels of selected transporter-encoding genes in roots during the Zn starvation and re-supply time series. A quadratic equation with one unknown y= ax2 + bx + c was used for the fitting, the parameters are listed in each graph. As tissues for elemental analysis and transcript profiling were obtained in independent experiments, each biological replicate where transcript levels were measured, was plotted against a randomly selected biological replicate from the elemental analysis dataset for the same time point. Time points post re-supply are depicted in shades of grey from –Zn (white) to +Zn (black).)

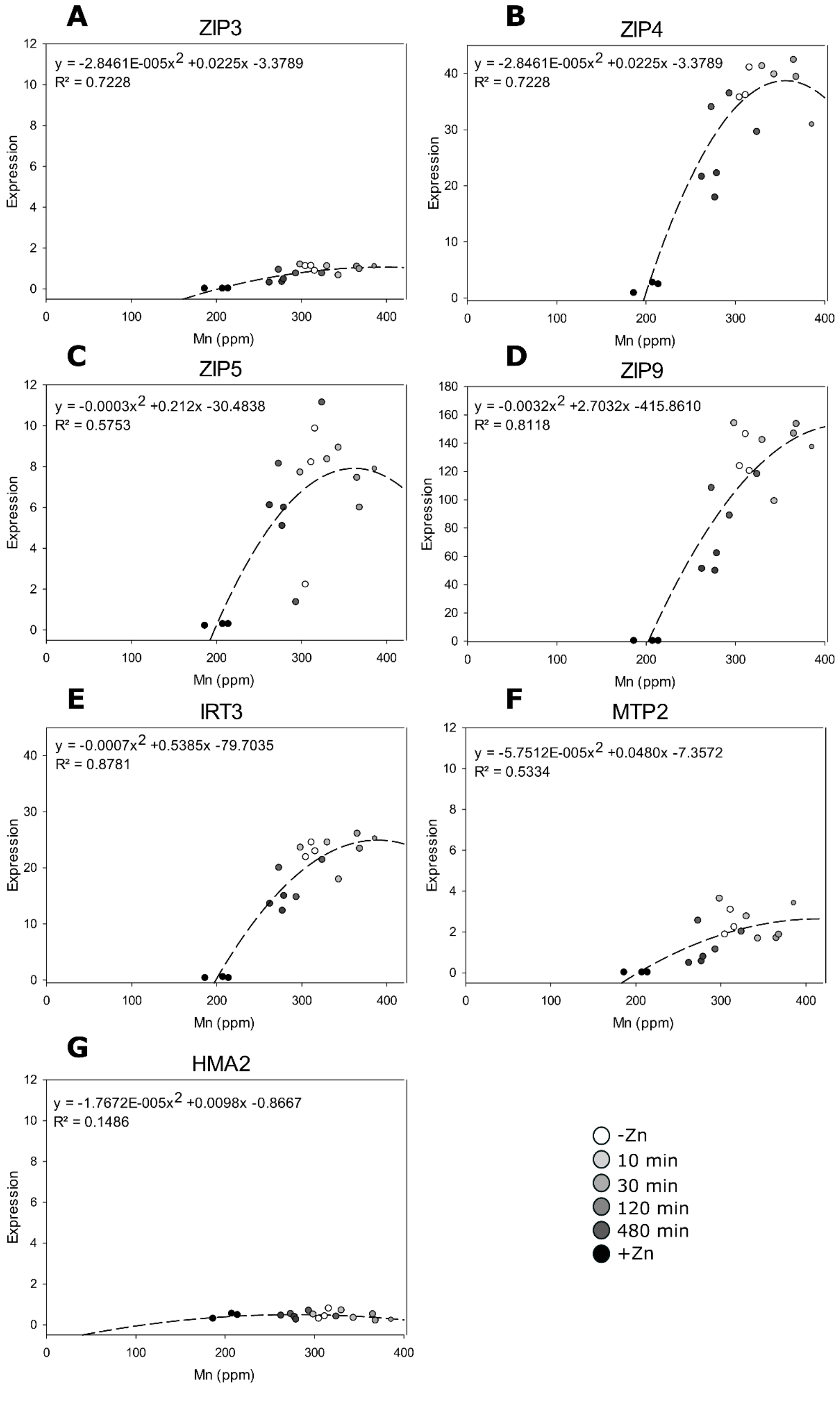


Figure S6. Relation between Mn levels and transcript levels of selected transporter-encoding genes in shoots during the Zn starvation and re-supply time series. A quadratic equation with one unknown y= ax2 + bx + c was used for the fitting, the parameters are listed in each graph. As tissues for elemental analysis and transcript profiling were obtained in independent experiments, each biological replicate where transcript levels were measured, was plotted against a randomly selected biological replicate from the elemental analysis dataset for the same time point. Time points post re-supply are depicted in shades of grey from –Zn (white) to +Zn (black).

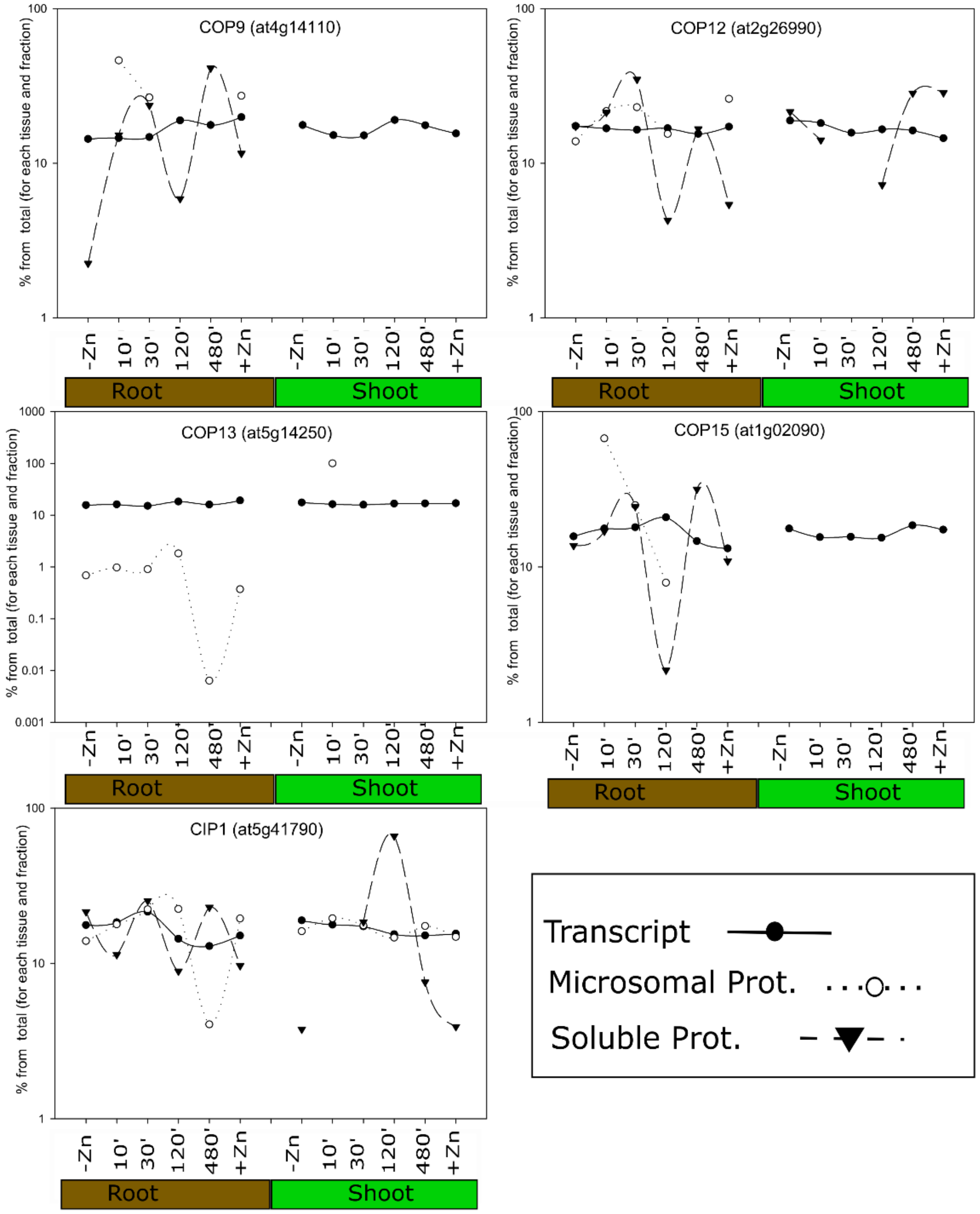


Figure S7. Transcript and protein levels of the COP 9 signalosome members. To present both transcript and protein levels for a gene on the same scale, the transcript expression/protein intensity is presented as the percent for the respective time point from the total of all time points in that tissue and fraction. The reader is reminded that these are relative values to start with and that transcript and protein intensities from each protein fraction are obtained in separate extraction steps. Relative protein intensity was obtained from cRacker (Zauber and Schulze, 2012), transcript levels are relative to *EF1α* and *At1g58050*, the value on the y-axis is common log, brown x-axis labels: root data, green x-axis labels: shoot data.

### Supplemental tables

Table S1: Sequence of primers for quantitative RT-PCR and T-DNA screening

| Transctipt name (AGI) | Forward primer | Reverse primer | Ref |
|---|---|---|---|
| ZIP3 (at2g32270) | GGAGTTTTGAACGCTGCATCC | CTGCGAGAAGGTCAACCAAAGA | Talke, I. N., et al. 2006 |
| ZIP4 (at1g10970) | AGCAAGAGAGGAATCAGGCTGC | CCAACCACAGGAACAACAGCA | Talke, I. N., et al. 2006 |
| ZIP5 (at1g05300) | CGTCACCGTGTCTGAAAGGA | CGTGGCAAAAGAGTCAACGG | |
| ZIP9 (at4g33020) | CCATCACTACTCCGATCGGTGT | CACCAATGCTGCAACGCTATAA | Talke, I. N., et al. 2006 |
| HMA2 (at4g30110) | TTGTCGCTGGCATTTATCCG | CAATTGTTGCTCCCACGGTT | |
| HMA5 (at1g63440) | GGTATCGACTCTGTTATCGCA | CCCGCAGCCTGTAATTCCTT | |
| MTP2 (at3g61940) | GCTTGGACATGATCACGGAC | CTCCAACAACTGCTCAGCTC | |
| MTP3 (at3g58810) | TTTGCCTTCCATCCCGAA | CTGATGTTGCAGCCTTTGCA | Haydon, M. J., et al. 2012 |
| IRT1 (at4g19690) | CCCCGCAAATGATGTTACCTT | GGTATCGCAAGAGCTGTGCAT | Becher, M., et al. 2004 |
| IRT3 (at1g60960) | AGTCATCCTCCTGGTCATGATT | GAGCATGACCAATGTCGAT | Talke, I. N., et al. 2006 |
| FRO2 (at1g01580) | ATTTACCCGATCGACCACAACA | TGCTCACGGAGATGACCAAGA | Becher, M., et al., 2004 |
| YSL6 (at3g27020) | TCACAGACTGGAGCCTAGCA | CGCAATGACACCACCATCAC | |
| NAS4 (at1g56430) | ACGACCAACTCGTAAACAAG | GAGAGTGTCGACATCTTCAC | Talke, I. N., et al. 2006 |
| bZIP19 (at4g35040) | TTCTCCCGGATGAGAGCGATGA | GCTGATTCACCGCCCTAAGCCT | Talke, I. N., et al. 2006 |
| bZIP23 (at2g16770) | GCTTCGTTGGAGGATGAGGT | GCTTCCAATGCAGCTTGACC | |
| VIT (at4g27870) | GGTGCCCCTCTTCTGTTTGA | CCATCACACCTGTTACTTTGGT | |
| TF (at1g02080) | GTTTCTCGGACCTGTGCTGA | TTGGCGTGGGAAGTTCAGTT | |
| COP9 (at4g14110) | ATGACGGAGGATGATGCCAC | GCTTGTTTCTTCACACTCGCC | |
| COP12 (at2g26990) | GGAAGAAGCAGCGACAGACT | ACTCCATCAGCATATTCGCCA | |
| COP15 (at1g02090) | TGGGCGGACAATATGAGTGAG | TCATGGACAGAGACTTCTTCAC | |
| COP13 (at5g14250) | AAGCGTTTGACTGCATTGC | TTGGCTGGTCAATCTCCACG | |
| CIP1 (at5g41790) | CGACACTGGAGCTGGAACTA | TTTTGCGCTTCCAACTGCTC | |

Table S2: Table S2. Enrichment of MapMan functional categories among the responding proteins in roots. Fisher's exact test, for a time point, was calculated using as background all possible protein comparison of the respective consecutive time-points and Zn deficiency and specified time point (from Data S5, as depicted in Fig. S4) for root tissues. The p-value was corrected for multiple comparisons (Benjamini-Hochberg). Full results are listed in Data S6. Enrichment of the entire functional bin is indicated by '*' and colored background, enrichment of one or more sub-bin is indicated by "sb" and colored text. Red font: proteins decreasing with time; blue font: proteins increasing with time; Bin: MapMan functional categories.

| | Root | 10 min | | 30 min | | 120 min | | 480 min | | +Zn | |
|---|---|---|---|---|---|---|---|---|---|---|---|
| BIN-CODE | Map Man Bin | 10min Root decreasing with time | 10min Root increasing with time | 30min Root decreasing with time | 30min Root increasing with time | 120min Root decreasing with time | 120min Root increasing with time | 480min Root decreasing with time | 480min Root increasing with time | +Zn Root decreasing with time | +Zn Root increasing with time |
| 1 | PS | | 1$^{sb}$ | 2$^{sb}$ | | 4$^{sb}$ | 1$^{sb}$ | 7$^{sb}$ | 3 | 3$^{sb}$ | 5$^{sb}$ |
| 2 | major CHO metabolism | | | | | 3 | | | 5$^{sb}$ | 3$^{sb}$ | 2 |
| 3 | minor CHO metabolism | 1$^{sb}$ | | | | 2 | | 4 | 1 | 1 | |
| 4 | glycolysis | | 1$^{sb}$ | | 3$^{sb}$ | 3 | | 5 | 9* | | 2 |
| 5 | fermentation | | | | 1 | | 1 | 4$^{sb}$ | 2 | | |
| 6 | gluconeogenesis / glyoxylate cycle | | | | | | | 1 | | | 1 |
| 7 | OPP | | 1$^{sb}$ | | 1 | 1 | | 5 | 4$^{sb}$ | | 4* |
| 8 | TCA / org transformation | | | | 2 | 7$^{sb}$ | | 13* | 12* | 2 | 7* |
| 9 | mitochondrial electron transport / ATP synthesis | | | 3$^{sb}$ | | 1* | 1 | 16$^{sb}$ | 2* | 5$^{sb}$ | 4 |
| 10 | cell wall | 1$^{sb}$ | 1 | 1 | 1 | 13$^{sb}$ | | 17$^{sb}$ | 16* | 3$^{sb}$ | 6 |
| 11 | lipid metabolism | 1 | | | 3 | 4* | 3$^{sb}$ | 21$^{sb}$ | 6 | 3 | 7$^{sb}$ |
| 12 | N-metabolism | | | | | | | 4$^{sb}$ | 5* | | 4* |
| 13 | amino acid metabolism | 2$^{sb}$ | 1$^{sb}$ | 1 | 4$^{sb}$ | 12 | 4$^{sb}$ | 15$^{sb}$ | 22$^{sb}$ | 5 | 10 |
| 14 | S-assimilation | 1$^{sb}$ | | | 1 | 1 | | 1 | 3 | 1 | |
| 15 | metal handling | | 3* | | | 4 | | 4 | 3 | 1 | 4* |
| 16 | secondary metabolism | | | | 5 | 23* | 2$^{sb}$ | 19$^{sb}$ | 19* | 8 | 12$^{sb}$ |
| 17 | hormone metabolism | 1$^{sb}$ | | 1$^{sb}$ | 1 | 9 | 4$^{sb}$ | 11$^{sb}$ | 10 | 3 | 4 |
| 18 | Co-factor and vitamine metabolism | | 1$^{sb}$ | 1$^{sb}$ | 1$^{sb}$ | 2 | | 3 | 1 | 1$^{sb}$ | |
| 19 | tetrapyrrole synthesis | 1$^{sb}$ | | | 1 | 2 | | 1 | | | |
| 20 | stress | 1 | 2$^{sb}$ | 3$^{sb}$ | 8$^{sb}$ | 16 | 9* | 30$^{sb}$ | 25$^{sb}$ | 22* | 11 |
| 21 | redox | | | | 2 | 6 | 1 | 11 | 11 | 1 | 8$^{sb}$ |
| 22 | polyamine metabolism | | | | | | | | | 1 | |

| | | | | | | | | | | | |
|---|---|---|---|---|---|---|---|---|---|---|---|
| 23 | nucleotide metabolism | | 1 | | 2[sb] | 7 | 1[sb] | 4* | 6 | 1 | 1 |
| 24 | Biodegradation of Xenobiotics | | | | 1 | 1 | | | 1 | | |
| 25 | C1-metabolism | | | 2* | | 1 | | 4 | 4 | 1[sb] | 1 |
| 26 | misc | 6* | 2 | 8[sb] | 6 | 41* | 22* | 68* | 42[sb] | 21[sb] | 26[sb] |
| 27 | RNA | 5* | | 3 | 12* | 34* | | 32[sb] | 31* | 19* | 11[sb] |
| 28 | DNA | | 1[sb] | 1 | 1 | 8 | 3[sb] | 17* | 4 | 4 | 8[sb] |
| 29 | protein | 6[sb] | 3[sb] | 8[sb] | 22* | 90[sb] | 2* | 104[sb] | 87[sb] | 33[sb] | 43[sb] |
| 30 | signalling | 1 | 4[sb] | 3 | 7[sb] | 17[sb] | 3[sb] | 41[sb] | 12* | 9 | 23[sb] |
| 31 | cell | | 3[sb] | 5[sb] | 7 | 20 | 3 | 22[sb] | 21 | 7 | 18 |
| 32 | micro RNA, natural antisense etc | | | | | | | | | | |
| 33 | development | 1 | | 2 | 8* | 9 | 4 | 19[sb] | 10 | 7 | 8 |
| 34 | transport | | 3[sb] | 1 | 7[sb] | 7* | 2 | 27 | 2* | 4[sb] | 13 |
| 35 | not assigned | 6 | 6 | 12 | 23 | 58* | 10 | 100[sb] | 44* | 36[sb] | 53 |

Table S3. Enrichment of MapMan functional categories among the responding proteins in roots. Fisher's exact test, for a time point, was calculated using as background all possible protein comparison of the respective consecutive time-points and Zn deficiency and specified time point (from Data S5, as depicted in Fig. S4) for shoot tissues. The p-value was corrected for multiple comparisons (Benjamini-Hochberg). Full results are listed in Data S6. Enrichment of the entire functional bin is indicated by '*' and colored background, enrichment of one or more sub-bin is indicated by "sb" and colored text. Red font: proteins decreasing with time; blue font: proteins increasing with time; Bin: MapMan functional categories.

| | Leaf | 10 min | | 30 min | | 120 min | | 480 min | | +Zn | |
|---|---|---|---|---|---|---|---|---|---|---|---|
| BIN-CODE | Map Man Bin | 10min Root decreasing with time | 10min Root increasing with time | 30min Root decreasing with time | 30min Root increasing with time | 120min Root decreasing with time | 120min Root increasing with time | 480min Root decreasing with time | 480min Root increasing with time | +Zn Root decreasing with time | +Zn Root increasing with time |
| 1 | PS | 4* | | 1 | 1 | | $4^{sb}$ | $2^{sb}$ | | | $3^{sb}$ |
| 2 | major CHO metabolism | | $1^{sb}$ | 1 | | | 3 | | | 4 | $1^{sb}$ |
| 3 | minor CHO metabolism | | | | | 1 | 1 | | 1 | $5^{sb}$ | 1 |
| 4 | glycolysis | | | | | | | | | 2 | $4^{sb}$ |
| 5 | fermentation | | | | | $1^{sb}$ | 1 | | $1^{sb}$ | 1 | |
| 6 | gluconeogenesis / glyoxylate cycle | | | | | | | | | 1 | |
| 7 | OPP | | | 1 | | | | | $1^{sb}$ | | |
| 8 | TCA / org transformation | | $1^{sb}$ | $1^{sb}$ | | | | | | 17* | 7* |
| 9 | mitochondrial electron transport / ATP synthesis | 4* | $2^{sb}$ | | 3* | 1 | 1 | | | $3^{sb}$ | |
| 10 | cell wall | | $1^{sb}$ | | $1^{sb}$ | | $3^{sb}$ | 1 | 1 | 3 | 8* |
| 11 | lipid metabolism | | $4^{sb}$ | | | | $4^{sb}$ | | | $14^{sb}$ | $8^{sb}$ |
| 12 | N-metabolism | | | 1 | | | | | | 2 | |
| 13 | amino acid metabolism | $1^{sb}$ | $2^{sb}$ | $2^{sb}$ | $1^{sb}$ | 1 | | | $1^{sb}$ | $7^{sb}$ | $6^{sb}$ |
| 14 | S-assimilation | | | | | $1^{sb}$ | | | | | 2* |
| 15 | metal handling | | 1 | 1 | | | | | | 7* | 3* |
| 16 | secondary metabolism | | $3^{sb}$ | $2^{sb}$ | | $1^{sb}$ | $1^{sb}$ | $2^{sb}$ | $3^{sb}$ | 8 | 14* |
| 17 | hormone metabolism | $1^{sb}$ | $2^{sb}$ | | $1^{sb}$ | $1^{sb}$ | 1 | | 2 | $7^{sb}$ | $3^{sb}$ |
| 18 | Co-factor and vitamine metabolism | | | | | | | 1 | 1 | | $2^{sb}$ |
| 19 | tetrapyrrole synthesis | $1^{sb}$ | $1^{sb}$ | $1^{sb}$ | | $1^{sb}$ | $2^{sb}$ | $1^{sb}$ | | | 6* |
| 20 | stress | 2 | $4^{sb}$ | 4 | | $3^{sb}$ | $6^{sb}$ | | 2 | $16^{sb}$ | $10^{sb}$ |
| 21 | redox | 1 | $1^{sb}$ | 1 | $1^{sb}$ | | 1 | | $1^{sb}$ | $12^{sb}$ | |
| 22 | polyamine metabolism | | | $1^{sb}$ | | | | | | | |
| 23 | nucleotide metabolism | 2 | 1 | $1^{sb}$ | | | | | | $4^{sb}$ | 1 |

| | | | | | | | | | | | |
|---|---|---|---|---|---|---|---|---|---|---|---|
| 24 | Biodegradation of Xenobiotics | | 1 | | | | | | 1[sb] | | |
| 25 | C1-metabolism | | | | | | 1 | | | | 2[sb] |
| 26 | misc | 1[sb] | 2 | 3 | 1 | 2 | 4 | 3 | 2[sb] | 40* | 11[sb] |
| 27 | RNA | 2 | 3[sb] | 2[sb] | 2 | 4[sb] | 7 | 2 | 4[sb] | 7 | 9[sb] |
| 28 | DNA | 2[sb] | | 2 | 2[sb] | 3* | 1 | | 2[sb] | 2 | 2 |
| 29 | protein | 4[sb] | 6[sb] | 12[sb] | 9[sb] | 7[sb] | 29[sb] | 11[sb] | 13[sb] | 37* | 26* |
| 30 | signalling | 3[sb] | 2 | 2[sb] | 2 | 2 | 11[sb] | 3[sb] | 2 | 24* | 7 |
| 31 | cell | | 1 | 2 | 3 | 6* | 2 | 2 | 2 | 9 | 9[sb] |
| 32 | micro RNA, natural antisense etc | | | | | | | | | | |
| 33 | development | | | 1 | | | 4 | 1 | | 6 | 2 |
| 34 | transport | 1 | 3 | 1 | 2 | 1 | 7[sb] | 1 | 1 | 15[sb] | 6[sb] |
| 35 | not assigned | 11 | 8 | 16 | 7 | 8[sb] | 30 | 6 | 10[sb] | 62[sb] | 29 |
